# Orthology inference at scale with FastOMA

**DOI:** 10.1101/2024.01.29.577392

**Authors:** Sina Majidian, Yannis Nevers, Ali Yazdizadeh Kharrazi, Alex Warwick Vesztrocy, Stefano Pascarelli, David Moi, Natasha Glover, Adrian M Altenhoff, Christophe Dessimoz

## Abstract

The surge in genome data, with ongoing efforts aiming to sequence 1.5M eukaryotes in a decade, could revolutionise genomics, revealing the origins, evolution, and genetic innovations of biological processes. Yet, traditional genomics methods scale poorly with such large datasets. Addressing this, “FastOMA” provides linear scalability, enabling the processing of thousands of eukaryotic genomes within a day. FastOMA maintains the high accuracy and resolution of the well-established OMA approach in benchmarks. FastOMA is available at https://github.com/DessimozLab/FastOMA/.

## Main

Within the decade, the Earth BioGenome initiative aims to sequence 1.5M eukaryotes^1^. This paves the way for understanding how all species evolved from life’s common origin. Yet due to processing limitations, even the thousands of genomes we have access to today are only studied piecemeal in practice. A fundamental step to comparative genomics analyses is to identify orthologs, genes of common ancestry that originated by speciation events^2^. When performed systematically, orthology delineation conveys how sequences were gained, lost or duplicated, assuming that their basic mode of inheritance is vertical descent. Deriving orthology allows for many types of downstream analysis, such as annotation propagation, phylogenomics, or phylogenetic profiling^3^.

State-of-the-art orthology methods face acute scalability issues^4^. Methods relying on all-against-all sequence comparisons can no longer keep up with today’s data, let alone tomorrow’s. For state-of-the-art pipelines such as our own OMA algorithm and database^5,6^, this translates to >10 million CPU hours to derive the orthology relationships of >2000 genomes that have been processed thus far. Methods relying on whole genome alignment, such as TOGA^7^, are more efficient but the genome alignment requirement limits their applicability to relatively closely related species. While “small-scale” comparative genomics has achieved remarkable progress, a more integrated, large-scale approach would be transformative.

To address this challenge, we introduce FastOMA, which dramatically speeds up orthology inference without sacrificing accuracy or resolution.

FastOMA is a complete rewrite of the OMA algorithm focused on scalability from the ground up **(Figure 1)**. By combining ultrafast homology clustering using k-mers, taxonomy-guided subsampling, and a highly efficient parallel computing approach, it achieves linear performance in the number of input genomes. First, we leverage our current knowledge of the sequence universe (with its evolutionary information stored in the OMA database) to efficiently place new sequences into coarse-grained families (Hierarchical Orthologous Groups ‘HOGs’ at the root level) using the alignment-free k-mer-based OMAmer tool^8^. In an attempt to detect homology among unplaced sequences (which could belong to families which are absent from our reference database), we then perform a round of clustering using the highly scalable Linclust software^9^. Next, we resolve the nested structure of the HOGs (Supplementary Information S1) corresponding to each ancestor, in an efficient leaf-to-root traversal of the species tree. By avoiding sequence comparisons across different families, the number of computations is drastically reduced compared to conventional approaches (see Online Methods for details).

**Figure 1.**
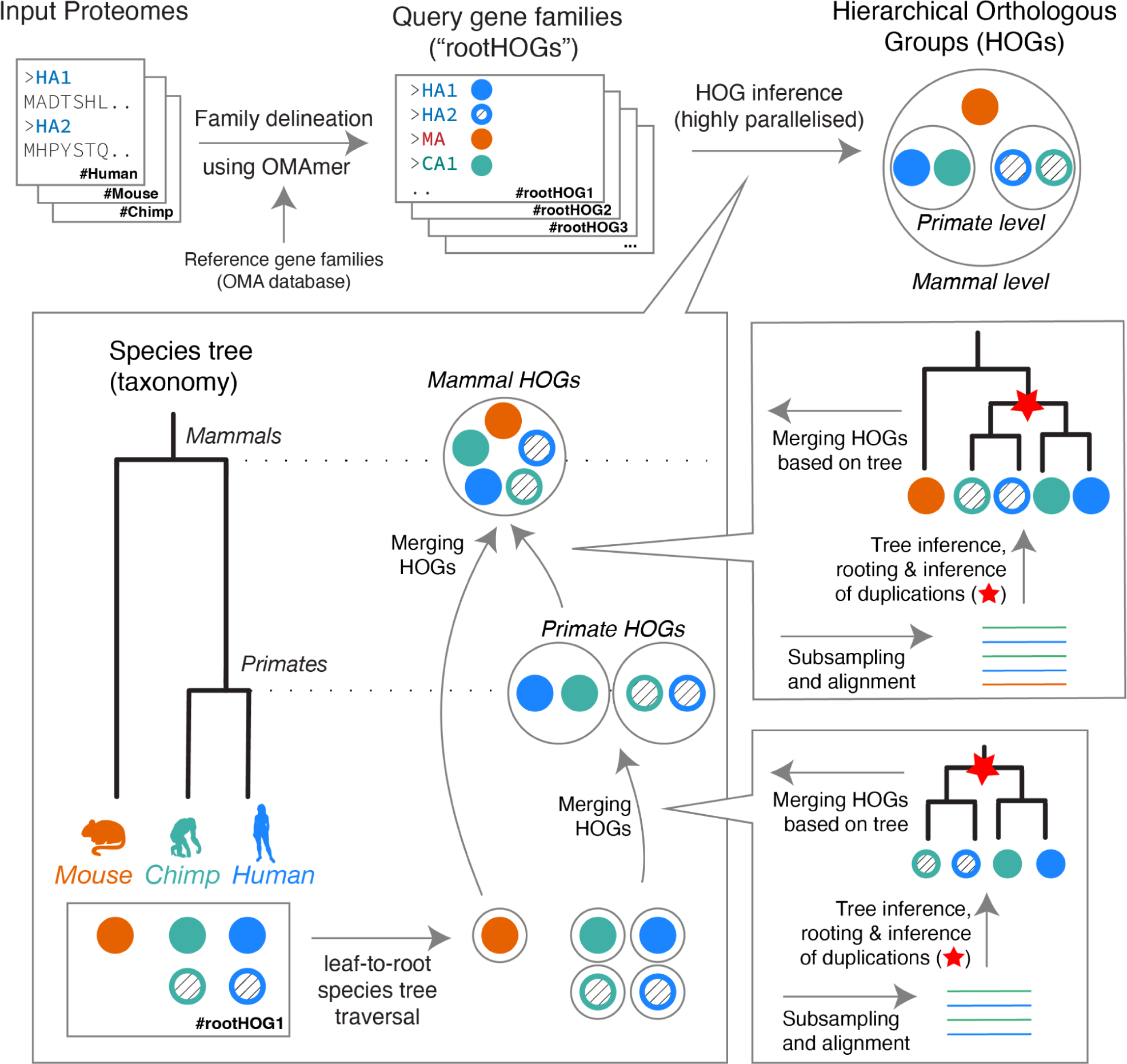
FastOMA algorithm overview. Input proteomes are mapped to reference gene families using the OMAmer software. HOGs are inferred using a “bottom-up” approach, starting from the leaves of the species tree and moving towards the root. At each taxonomic level, HOGs from the child level are merged, resulting in HOGs at the current level. To decide which HOGs should be merged, sequences from the child HOGs are used to create a multiple sequence alignment, followed by gene tree inference to identify speciation and duplication events. Child HOGs are merged if their genes evolved through speciation (see **Methods** section & **Supplementary Information** for details).

FastOMA has unprecedented scalability without sacrificing accuracy in a diverse range of benchmarks. We assessed the accuracy of FastOMA on the Quest for Orthologs suite of benchmarks^10^. FastOMA retains OMA’s high precision accuracy, and even improves upon it in terms of recall, positioning it on the Pareto frontier of orthology inference methods. For instance, on the SwissTree reference gene phylogeny benchmark, FastOMA outperforms other methods with a precision of 0.955 in reference gene phylogenies (**Figure 2a**). With a recall in line with most state-of-the-art methods (0.69, lower than those of Panther and OrthoFinder), the balance of these metrics indicates a well-tuned approach to orthology inference, with a focus on minimising false positives. Likewise, on the Generalised Species Tree benchmark at the Eukaryota level, FastOMA is amongst those with the lowest topological error, with a normalised Robinson-Foulds distance – the number of edges in common between two trees normalised by the total number of edges– of 0.225 to the reference tree, at moderate recall (**Figure 2b** and Supplementary Information S2-8).

**Figure 2.**
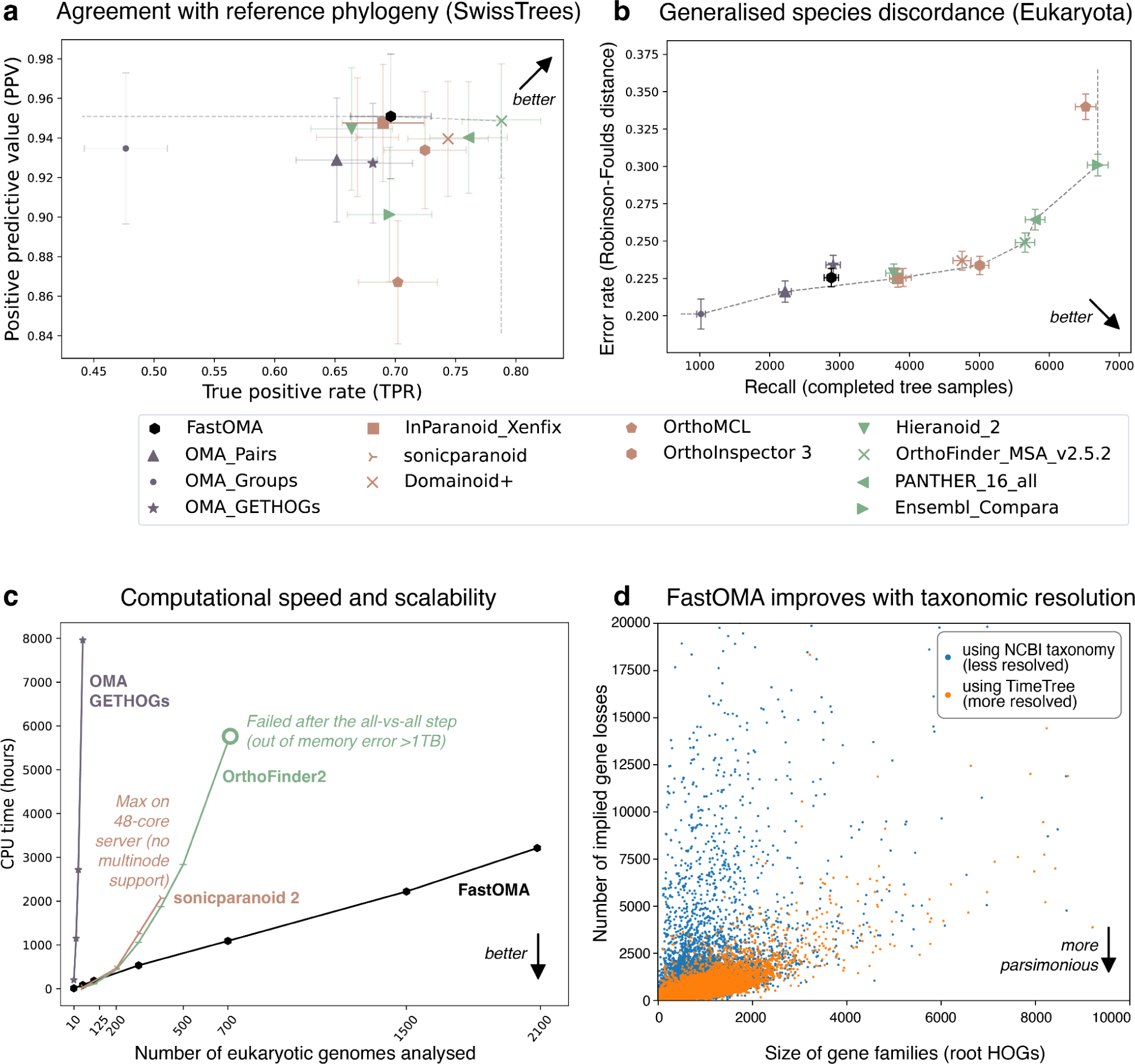
FastOMA is not only fast but also accurate: a) Quest for Orthologs (QfO) benchmark, agreement with SwissTree reference phylogeny; b) QfO benchmarking of generalised species discordance test on Eukaryota clade, where the gene tree inferred from orthologous genes are compared with the reference species tree (see **Supplementary Information S2.1** for detail); c) computation time comparison of FastOMA and state-of-the-art alternatives; d) impact of species tree resolution on the complexity of gene family evolutionary scenario (proxied by the number of gene losses over the gene family history). Each point represents a gene family (a “root HOG”), whereby the size of a gene family corresponds to the number of genes in it. (The figure is truncated to focus on the most relevant region; a version with all data in **Supplementary Figure 24**; See Methods for implied losses calculation).

A key achievement of FastOMA is its linear scaling behaviour (**Figure 2c**), which opens up the possibility of processing extensive datasets rapidly. FastOMA inferred orthology among all 2,086 eukaryotic UniProt reference proteomes in under 24 hours, using 300 CPU cores. In the same timespan, the original OMA algorithm could only process 50 genomes. Even methods optimised for speed such as OrthoFinder^9^ or SonicParanoid^10^ still exhibit quadratic time complexity (**Figure 2c**). Thus, FastOMA’s linear scalability breaks new ground.

The initial sequence placement step using OMAmer helps FastOMA achieve its speed, but the subsequent alignment and tree inference steps are critical for its accuracy. Indeed, sequence placement alone is considerably less accurate than state-of-the-art methods in benchmarks (Supplementary Information S3).

FastOMA exploits known taxonomic relationships to reduce the number of sequence comparisons. By default, it relies on the commonly used NCBI taxonomy^11^, but users can specify any reference species phylogeny as input. To assess the impact of the resolution of the input tree on orthology accuracy, we compared FastOMA’s performance on UniProt reference proteomes with a more resolved species tree derived from the TimeTree resource^12^. Compared with the NCBI taxonomy, this resulted in improved ortholog predictions, with more parsimonious gene family evolution history, lowering the number of implied gene losses across all gene families (**Figure 2d**). FastOMA is also robust to errors artificially introduced in the species taxonomy (**Supplementary Figure 18-20**). FastOMA can thus use advances in taxonomic knowledge for better orthology predictions and will benefit from the higher resolution that is brought by new genomic sequences from large-scale sequencing projects.

FastOMA contains additional features that make it easier to deal with complex and noisy genomic data. It is designed to handle multiple isoforms for the genes resulting from alternative splicing and select the most evolutionarily conserved ones, and can also deal with fragmented gene models. Both features lead to noticeable improvements in FastOMA inferences (Supplementary Information S9-10). As it uses the same data structure as OMA, FastOMA benefits from its rich ecosystem of downstream applications which includes phylogenetic profiling, efficient gene family visualisation, ancestral synteny inference, and advanced phylostratigraphy, enabling researchers to trace gene family histories and understand gene emergence, duplication, and loss events^5,13^

In conclusion, the FastOMA algorithm offers a unique solution for accurate orthology inference, making it possible to study evolutionary history at the scale of massive genomics projects.

## Online Methods

### FastOMA algorithm outline

FastOMA is a method for inferring orthology relationships. The input to FastOMA includes the proteome sets of species and the species tree. The FastOMA algorithm consists of two main steps: finding rootHOGs and inferring the nested structure of HOGs (**Figure 1**).

### Step 1) FastOMA gene family inference

The FastOMA algorithm infers gene families from the provided proteomes. The process begins by mapping the input proteomes onto the reference HOGs (Supplementary Information S1) using the OMAmer tool (**Figure 1**). Proteins mapped to the same reference HOG are then grouped together, forming query rootHOGs, with the exclusion of proteins already present in the database. Thus, proteins in the database reference HOGs are not used in the next steps in FastOMA.

While each rootHOG ideally represents a single gene family, instances may arise where a gene family of query proteomes is split among multiple rootHOGs. To address this, FastOMA tries to find those query rootHOGs that are associated with the same gene family. FastOMA leverages the ability of OMAmer to report multiple rootHOGs to which the sequences could be mapped, along with their score. This score (‘family_p’) is the p-value of having as many or more k-mers in common between the protein sequence and the HOG under a binomial distribution, reported in negative natural logarithm. Considering a minimum threshold of 70 (by default), we construct a graph of rootHOGs, where each node represents a query rootHOG. In such a graph, we add an edge between two nodes (rootHOGs) when a minimum of 10 proteins (by default) are mapped to both query rootHOGs and it represents at least either 80% of all proteins mapping to the bigger rootHOG or 90% of those mapping to the smallest one. This ensures a high overlap of protein content of the merged rootHOG. Finally, we group the members of all HOGs in each highly connected component^14^ of this graph in a single query rootHOG.

It is worth noting that some proteins may not be assigned to any reference HOGs due to no recognisable homologs in the reference database. Additionally, there is a scenario where only one protein is mapped to the rootHOG, referred to as a singleton, representing an individual rather than a group^5^. To ensure those genes are not lost to FastOMA’s orthology inference, these singletons and unmapped sequences are combined into a FASTA file on which we run Linclust, the clustering tool from the MMseqs package^9^. This yields new query rootHOGs.

Critically, assigning proteins to rootHOGs (gene families) allows us to avoid unnecessary all-against-all comparisons of unrelated proteins (those without homology), thanks to the speed of OMAmer and Linclust. All of the query rootHOGs are written as FASTA files to be used in the next step and can be handled in parallel.

Notably, the OMA team provides regular updates to the OMA database, increasing the number and diversity of species included in the database used by OMAmer. This results in higher resolution for k-mer based grouping. As more taxa get included, we foresee FastOMA’s inference will improve as more sequences are placed into rootHOGs.

### Step 2) FastOMA orthology inference

For every query rootHOG, FastOMA infers the nested structure of the HOG (as depicted in **Figure 1**). The objective is to identify the genes that are grouped together at each taxonomic level as a HOG; which means they descended from a single gene at that specific level. Note that the number of HOGs at each level reflects the number of copies of the gene present in the ancestral species.

To achieve this, FastOMA follows a bottom-up approach by traversing the species tree. Starting from the leaves of the tree (extant species), each gene in the species’ proteome is treated as a HOG. At each level in the traversal, certain HOGs from the child level are combined. The determination of which HOGs will be merged is guided by a gene tree containing the proteins of species descending from this node. The merging is done for all HOGs that descended from the same common ancestor by a speciation event. The entire process is detailed below:

#### Gene tree inference

All the proteins in HOGs at the child level are collectively used for generating a multiple sequence alignment (MSA) using the MAFFT package^15^. As part of the FastOMA python script, the MSA undergoes column-wise trimming with a default threshold of 0.2, meaning that we remove columns of the MSA that have more than 80% gap elements (see Supplementary Information S5). Those aligned sequences (rows in MSA) that exceed a default threshold of >50% gaps are subsequently removed. However, we keep them in the HOG but they are not used for tree inference. Subsequently, we employ FastTree^16^ to infer the gene tree, and this tree is rooted using the midpoint approach^17^.

To expedite the orthology inference process at deeper levels of the trees where the number of children is prohibitively high, we implement a subsampling approach, retaining only a specified number of proteins per HOG (see **Supplementary Figure 12-14**, by default 20 proteins are randomly selected) used for the multiple sequence alignment (MSA) and tree inference. And, the unsampled sequences will have the same fate as the rest of the proteins in the same group at the defined taxonomic level.

Note that the subsampling strategy is key to the speed of FastOMA, and expectedly there is a trade-off between accuracy and speed. Our benchmarking results indicate that FastOMA performs well with the subsampling approach, but users can change the degree of the subsampling in the parameter file.

#### Duplication and speciation event labelling

Each internal node in the gene tree is classified as either a duplication or a speciation event using the species overlap method^18^. For each node in the gene tree, this involves calculating the ratio of the number of shared species between its two subtrees divided by the number of all species (union). If the ratio equals zero, the node is labelled as a speciation event; otherwise, it is labelled as a duplication event. When the species overlap ratio is less than 0.1 (as per default settings), indicating very low support for a duplication event, all leaves from the child subtree with the least number of proteins are excluded from merging decisions (described in the next subsection *“HOG merging”*). In other words, these proteins will stay in the corresponding HOGs as it was in the previous taxonomic level, and only the taxonomic label of the HOG is updated to the current taxonomic level (assuming no other merging happens in another part of the gene tree for this HOG). This is done to ensure that errors in gene annotation or inaccurate tree inference minimally affect the orthology inference.

#### HOG merging

Starting from the root of the gene tree, evidence of a speciation event (i.e., the internal node annotated as a speciation event due to no species overlap) prompts the merging of the HOGs of the leaves descending from the nodes. This is achieved by constructing a HOG graph, where each node represents a HOG. An edge is introduced between HOG1 and HOG2 if protein1 (located in HOG1) and protein2 (in HOG2) coalesce at a speciation event in the gene tree. Subsequently, each connected component within this graph constitutes a HOG at the current level of the species tree. Furthermore, FastOMA has a mechanism to handle spuriously merged subHOGs; at the deeper taxonomy level, when genes within a subHOG coalesce at a duplication event in the gene tree, FastOMA splits the subHOG into two, ensuring copies of ancestral genes are not co-present in a subHOG.

#### Inferring orthology relationship

Once the species tree traversal is complete, the nested structure of the query HOG is fully resolved. From the HOG structure inferred this way, all orthology and paralogy relationships can be efficiently deduced.

#### Note on parallelisation

Scalability has been a major challenge in the field of orthology inference highlighted by the Quest for Orthologs community for many years^10,27^. FastOMA is optimised to process taxonomic levels in parallel (when possible) by inferring HOGs at all taxonomic levels, accounting for dependencies among child HOGs– i.e., a node will be processed after all its child nodes are processed. To optimise parallelization efficiency by avoiding unnecessary overheads of Nextflow and Slurm management workflows, FastOMA groups approximately 150 small to medium-sized query rootHOGs together, treating them as a single job. Conversely, large rootHOGs are processed individually (to infer nested structure of HOGs) for optimal performance using python-future for which taxonomic parallelization is activated. The default rootHOG file size threshold for this purpose is 400k bytes, ∼500 proteins (See Supplementary Information S11).

### FastOMA outputs

The main output of FastOMA is an OrthoXML file which stores HOGs and their nested structures, allowing to reconstruct their evolutionary histories. Furthermore, FastOMA reports the protein list in each rootHOG (gene family) in TSV format. A final FastOMA output is a list of proteins in strict orthologous groups, wherein all genes within the group are orthologous to each other, which can be used as marker genes for phylogenetic analyses^19,20^. Besides, the user can store the gene trees and multiple sequence alignments of the subsampled HOGs for all taxonomic levels.

### Isoform selection

FastOMA is capable of handling proteomes that feature multiple protein isoforms for a gene due to alternative splicing. Users can provide an isoform file where each row lists comma-separated protein IDs associated with a gene. FastOMA selects the isoform with the highest ‘family_p’ score, the one with the best fit to known proteins in the reference rootHOG based on k-mer content. For the evaluation of isoform selection, we used the UniProt reference proteomes and their splice information https://ftp.uniprot.org/pub/databases/uniprot/current_release/knowledgebase/reference_proteomes/E ukaryota.

### FastOMA Software

The FastOMA codebase is composed of multiple subpackages written in Python. FastOMA benefits from the Nextflow workflow^21^ to parallelize different steps and subpackages considering the dependencies modelled as a direct acyclic graph (Supplementary Information S11). The software is publicly available at https://github.com/DessimozLab/FastOMA and on DockerHub https://hub.docker.com/r/dessimozlab/fastoma.

### Time comparison on Eukaryotic dataset

We considered all the 2181 eukaryotic UniProt reference proteomes (accessed on 25 January 2023), and filtered them to keep those with a minimum BUSCO Completeness of 50%, resulting in 2086 proteomes in total. We ran SonicParanoid^22^, OrthoFinder^23^, and FastOMA on datasets with different sizes ranging from 10 to 2086 species. OrthoFinder 2.5.4 was run in two steps. First, to generate all-against-all sequence comparisons, we used the -op parameter to generate and execute command lines for Diamond. Then, the rest of OrthoFinder was conducted. SonicParanoid 2.0.4 was used with default parameters using 48 CPUs with a limit of 3 days wall clock. It is neither possible to parallelize SonicParanoid2 on different computation nodes nor feed it with the result of Diamond, hence we could not obtain compute time for the larger datasets during the mentioned time limit. For FastOMA, the NCBI tree was used by downloading via the ETE3 package^24^. The comparison of tools in terms of wall-clock time in hours is reported in **Supplementary Figure 25**. The Diamond part of OrthoFinder and all steps of FastOMA use different nodes on the cluster, so the reported wall-clock time might have been affected by the availability of nodes at the time of each run. However, the CPU times reported in Figure 2c are more accurate.

### Analysis on tree resolution

We ran FastOMA on both the TimeTree and the NCBI tree. For the TimeTree analysis, we uploaded the list of species names to the TimeTree webserver^12^ (https://timetree.org). This resulted in a species tree with 1757 leaves since some of the species were not available in TimeTree. We ran FastOMA with default parameters on the dataset of 1757 proteomes and with both the TimeTree tree and NCBI tree as the species tree. We used pyHAM^25^ for calculating the implied gene losses.

To calculate the estimated proportion of proteomes composed of fragments, we ran OMArk^26^ v0.3 on all proteomes. We used the BUSCO statistics downloaded from the UniProt website for the full Eukaryotic dataset.

We also conducted another analysis to study the impact of the species tree for the QfO dataset where five pairs of species are swapped. The results are provided in **Supplementary Information S7** and **Supplementary Figures 18-20**, where FastOMA shows a moderate level of robustness. However, having an erroneous species tree impacted the orthology inference by introducing false positives.

To conclude, we highlight that the orthologous and paralogous genes are found using the species overlap method on the gene tree and the species tree is used to determine the order of comparisons, defining the HOG structure. Thus, a fully-resolved species tree is not needed to infer orthology information with FastOMA. However, errors in the species tree can potentially propagate through the orthology inference process.

### Benchmarking against the QfO reference proteome set

We ran FastOMA on the 78 reference proteomes used in the QfO benchmark and the associated standard species trees as input. We then submitted the results to the Quest for Orthologs benchmarking service^4,10,27^ and obtained the results on the 11 available benchmarks. In these benchmarks, FastOMA is compared to several state-of-the-art methods that are available in the QfO public resource including EnsemblCompara^28^, Domainoid^29^, OrthoMCL^30^, Ortholnspector^31^, sonicparanoid^32^, PANTHER^33^, OrthoFinder^23^, Hieranoid^34^ and the OMA family^35–38^. QfO analysis is described in detail in the Supplementary Information S2.

### Analysis of the QfO reference proteome set using InterProScan classification of protein families

To study the influence of OMA database and OMAmer on the performance of FastOMA, we replaced the first part of the procedure, normally done by placing query genes into the OMA database rootHOGs with OMAmer, with InterProScan. We used InterProScan to group the QfO proteomes into gene families predefined by InterProScan^39^. To do so, we first ran InterProScan with the argument *-appl Pfam* on the QfO dataset which grouped the proteins into InterProScan families^40^. Then, we created the rootHOG with those groups, maintaining the same InterProScan family identifier. Then, we ran the rest of FastOMA on these rootHOG FASTA files. The QfO benchmarking results are shown in **Supplementary Information S6** and **Supplementary Figures 15-17**. Note that a user can provide their own initial grouping of proteins to be used with FastOMA. This could be put in practice in two ways: 1) running the last two processes of *FastOMA.nf* (*hog_rest* and *collect_subhog*) on the user’s protein family in FASTA format. 2) Providing group mapping of proteins in the OMAmer format.

### Computations

All the analyses were conducted on the high-performance computer cluster of the University of Lausanne which houses 96 computation nodes. Each node is equipped with two 24-core AMD CPUs, totaling 48 cores per node. Data was written and read on a 150 TB SSD scratch drive. For the QfO analysis, most steps of FastOMA needed less than 10 GB of memory and it peaked at 32 GB.

## Data availability

UniProt reference proteomes and splice information (_additional.fasta.gz) were downloaded from https://ftp.uniprot.org/pub/databases/uniprot/current_release/knowledgebase/reference_proteomes/Eukaryota. The 2020 version of QfO proteomes was downloaded from the EBI repository at http://ftp.ebi.ac.uk/pub/databases/reference_proteomes/previous_releases/qfo_release-2020_04_with_updated_UP000008143/. The OMAmer database used in this study is available at https://omabrowser.org/All/LUCA-v2.0.0.h5. The OMAmer database, an archive of FastOMA code, the Time tree with annotation of internal nodes of 1757 species in Newick format, the UniProt IDs, and the inferred HOG for 1757 Eukaryotic species in OrthoXML format are all deposited at www.doi.org/10.5281/zenodo.10403053/.

## Code availability

FastOMA is free open-source software (Mozilla Public License 2.0) available at https://github.com/DessimozLab/FastOMA. We used the publicly available code for the QfO benchmarking test which is available at https://github.com/qfo/benchmark-webservice

## Acknowledgement

We thank Clement Train for updating PyHam.

## Author contributions

SM, AA and CD developed the method. SM and AA implemented the software. SM, AYK, YN, AW, NG, DM, SP contributed to the analysis. CD and SM wrote and edited the manuscript. All authors read and approved the final version of the manuscript.

## Supplementary Information

S1. A note on the definition of HOG

S2. Full benchmarking results for QfO

S3. Comparing FastOMA results with OMAmer mapping

S4. Impact of reference HOG database on FastOMA results S5. FastOMA robustness on threshold

S6. Impact of OMA database on FastOMA

S7. Impact of species tree on FastOMA

S8. The group benchmarking for the clade Bilateria.

S9. FastOMA’s ability to select isoforms

S10. FastOMA’s ability to find split genes

S11. FastOMA Nextflow DAG

## S1. A note on the definition of HOG

A HOG comprises all the present-day genes that have descended from a single gene in a reference ancestor^38^. Hence, HOGs relate present-day genes in terms of those of ancestral species. For instance, all mammalian insulin genes descended from a single insulin at the root of the mammals. There is thus one insulin HOG at the mammalian level. But within rodents, where insulin is duplicated, the two copies belong to distinct rodent HOGs, nested into the first one (**Supplementary Figure 1**).

HOGs have several conceptual advantages: HOGs provide a precise definition for the useful but vague concepts of gene families and subfamilies. Because each HOG corresponds to an ancestral gene in a given ancestor, they collectively give the gene repertoires of said ancestor. HOGs are a scalable alternative to gene trees, which tend to be hard to infer and interpret.

**Supplementary Figure 1.**
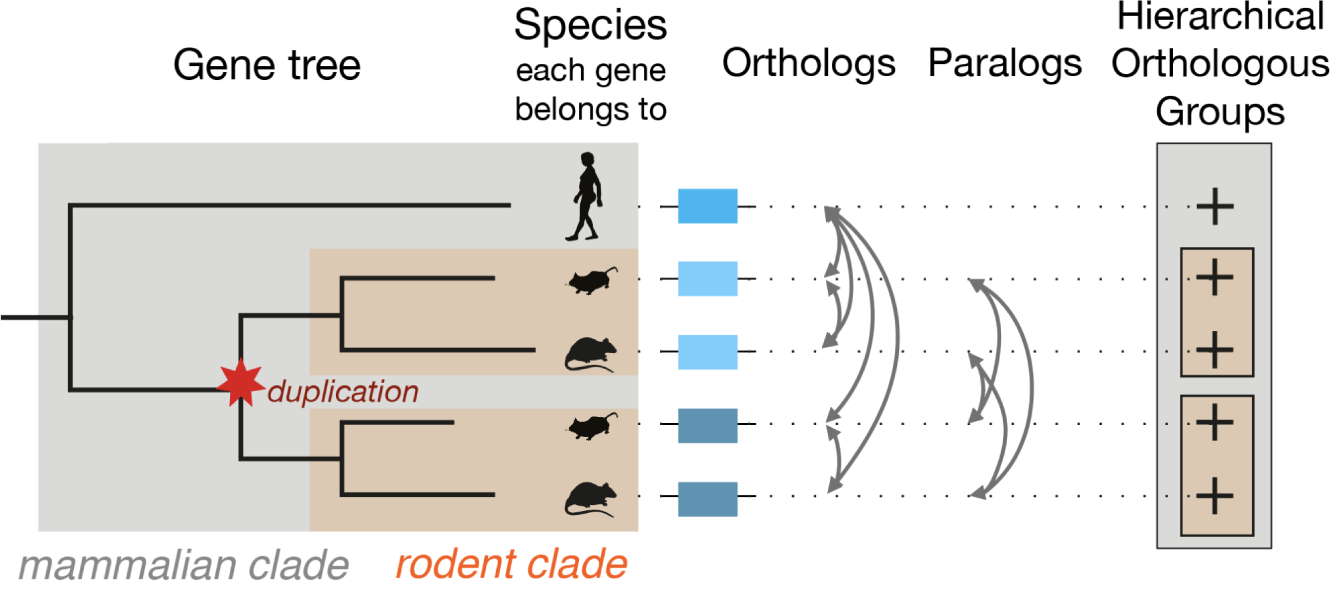
An example of HOG structure^41^.

The “rootHOG” corresponds to the deepest ancestral gene and defines a gene family. Of note, rootHOG may correspond to a specific taxonomic level of the species tree (not necessarily the root of the species tree) where it appeared first over the course of evolution. When multiple HOGs exist in a descendant, they represent distinct gene subfamilies. In a nutshell, HOG is the fundamental underlying evolutionary concept which ties in the concepts of present-day or ancestral genes (HOGs), gene families (rootHOGs), and gene subfamilies (child HOGs).

The standard for reporting HOGs (and most orthology inferences) is the orthoXML format (https://orthoxml.org/) which allows to describe nested orthologs and paralogs groups (**Supplementary Figure 2**). Thus, an orthoXML file is the primary output of FastOMA.

**Supplementary Figure 2.**
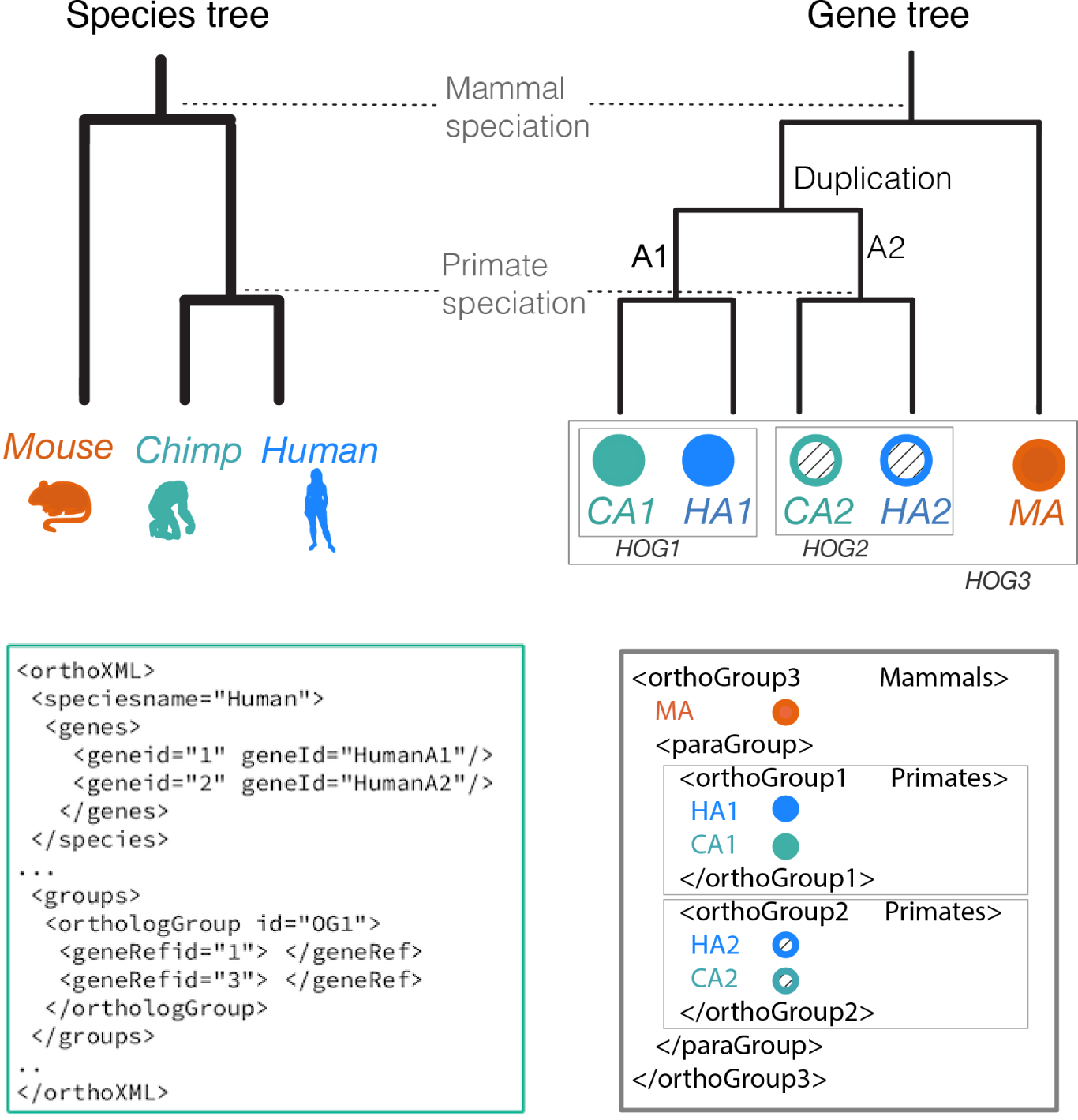
An example of HOG structure and orthoXML format.

## S2. Full benchmarking results for QfO

The accuracy of FastOMA was evaluated using the 2020 version of the Quest for Orthologs (QfO) benchmarking dataset^4,27^. This includes 78 species across the tree of life with 984,137 protein sequences for which state-of-the-art orthology inference methods were run. The QfO benchmarks are a series of 11 different tests in three categories including the species discordance test, agreement with reference phylogeny, and functional analysis which are presented below.

### S2.1 Species tree discordance test

As one usage of orthology is to infer species trees, as part of the QfO benchmarking, we conducted the species tree discordance test. This evaluates ortholog accuracy by assessing the accuracy of the species tree reconstructed based on it. To decrease the gene-species tree discrepancies due to incomplete lineage sorting, orthologs are sampled from species separated by more than 10 million years^42^. This test is designed for three clades including Eukaryota, Bacteria and Fungi. The results are provided in **Supplementary Figure 3**.

The subplots a-c in this figure are dedicated to the number of trees that were completed (using orthologous pairs as a proxy for recall) and for which Robinson-Foulds (RF) distance (to compare the topological differences between two trees) were calculated. We use the ETE3 package to calculate this metric. In principle, to calculate the RF distance between two trees, first, each tree is cut into two subtrees each time by removing a branch. Then, the sets of leaves in each partition are found and compared to the partitions of the partitions of the other tree. The number of different partitions is reported as RF distance. The value is normalised by dividing it by the total number of present splits.

In this benchmark (**Supplementary Figure 3a-c**), FastOMA performs well, with a low average Robinson-Foulds distance (higher precision) and moderate recall, as reflected in the number of completed tree samplings and number of orthologous pairs. This places FastOMA close to the Pareto frontier for three clades of Eukaryota, Fungi, and Bacteria.

The subplots **d-f** in **Supplementary Figure 3** show the same benchmark data, but use the fraction of incorrect trees (gene trees with at least one difference from the species tree) as a measure of accuracy and the number of orthologs as a measure of recall.

FastOMA results are similarly close to the Pareto frontier with this measure for Eukaryota and Fungi. It does not perform better than other methods in Bacteria, with the proportion of incorrect trees being higher than other methods with a similar number of orthologs. This might be related to the fact that the FastOMA algorithm for orthology calling assumes vertical descent between genes, an assumption that is not always met in Bacteria.

Since the number of species under study in this test is limited, the variance is high. This leads to the generalised species tree discordance test described below.

### S2.2 Generalised species tree discordance test

The generalised species tree discordance test (GSTD) includes Eukaryota, Vertebrata, Fungi and LUCA levels. The results reported in **Supplementary Figure 4** are measured in terms of number of orthologous pairs, number of completed tree sampling (as proxies for recall), Robinson-Fold distance, and the fraction of incorrect completed trees (as proxies for precision). In this test, OMA-groups has the highest precision and lowest recall; OrthoMCL and Ensembl Compara are at the other extreme with the highest recall and lowest precision overall. FastOMA consistently has a better recall than other OMA predictions with a higher RF distance compared to OMA-GETHOGs2. Over the benchmark of different clades, it ranks at or close to the Pareto frontier, between the other OMA predictions and most of the other included methods. The relative ranking varies between clades, with FastOMA having slightly lower recall in Vertebrates but higher sensitivity than some OMA predictions for example, but stays true to the general trend overall.

### S2.3 Reference gene phylogenies

Another orthology benchmark we exploited is based on the reference gene phylogenies^43^. We used SwissTree, which is a small collection of large- and high-confidence gene family phylogenies with different types of challenges for orthology prediction and species from all domains. In this benchmark, FastOMA performs comparably to other methods, with one of the highest precision (true positive rate: 0.95), but a moderate recall (positive predictive value: 0.69). We also calculated the test for the TreeFam-A reference gene phylogeny which is a larger set of metazoan gene trees covering a taxonomically restricted but wider range of protein families. In this benchmark, FastOMA ranks close to other OMA predictions, with a higher prediction than other tree-based approaches but lower precision and recall than other graph-based predictions. These are reported in **Supplementary Figure 5**.

### S2.4 Gene ontology conservation benchmark

The Gene Ontology (GO) conservation benchmark shows how well the Gene Ontology annotations are conserved among the predicted orthologs. This test is based on studies that have demonstrated that orthologs exhibit significant (but moderate) conservation in terms of GO annotation similarity as opposed to paralogs^10^. Therefore, accurate inference of orthology is expected to be associated with gene pairs that are functionally similar at a given evolutionary distance. We assessed functional similarity based on experimentally-backed annotations from the UniProt–Gene Ontology Annotation (GOA) database and Enzyme Commission (EC) numbers from the ENZYME database. To benchmark, we calculated the average Schlicker semantic similarity between GO and EC terms of predicted orthologous pairs as a measure of precision and the number of predicted ortholog relationships as recall^10^. The average Schlicker of FastOMA is 0.465 (0.925) in GO (EC), placing it close to the Pareto frontier.

**Supplementary Figure 3.**
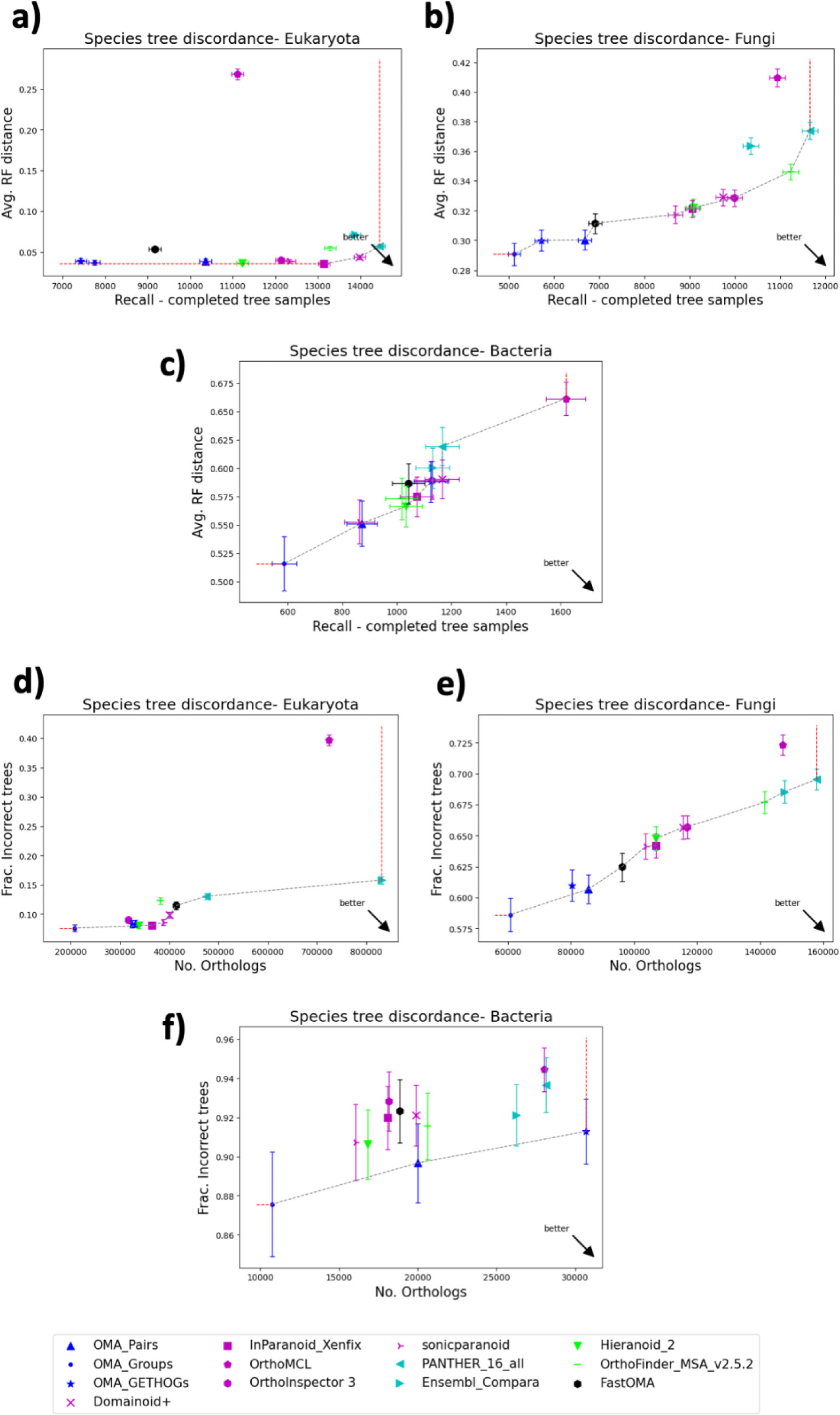
The result of the species tree discordance test. (a-c) in terms of average Robinson-Foulds distance vs number of completed tree samples (a-c) and in terms of fraction of incorrect trees and number of orthologs (d-f). The other methods of OMA are in blue and the new **FastOMA** is in black. Graph-based methods (OrthoMCL, ORthoInspector, InParanoid, Sonicparanoid, and Domainoid+) are in purple and the tree-based methods are in cyan. The hybrid methods which use both gene tree and graph structure are in green.

**Supplementary Figure 4.**
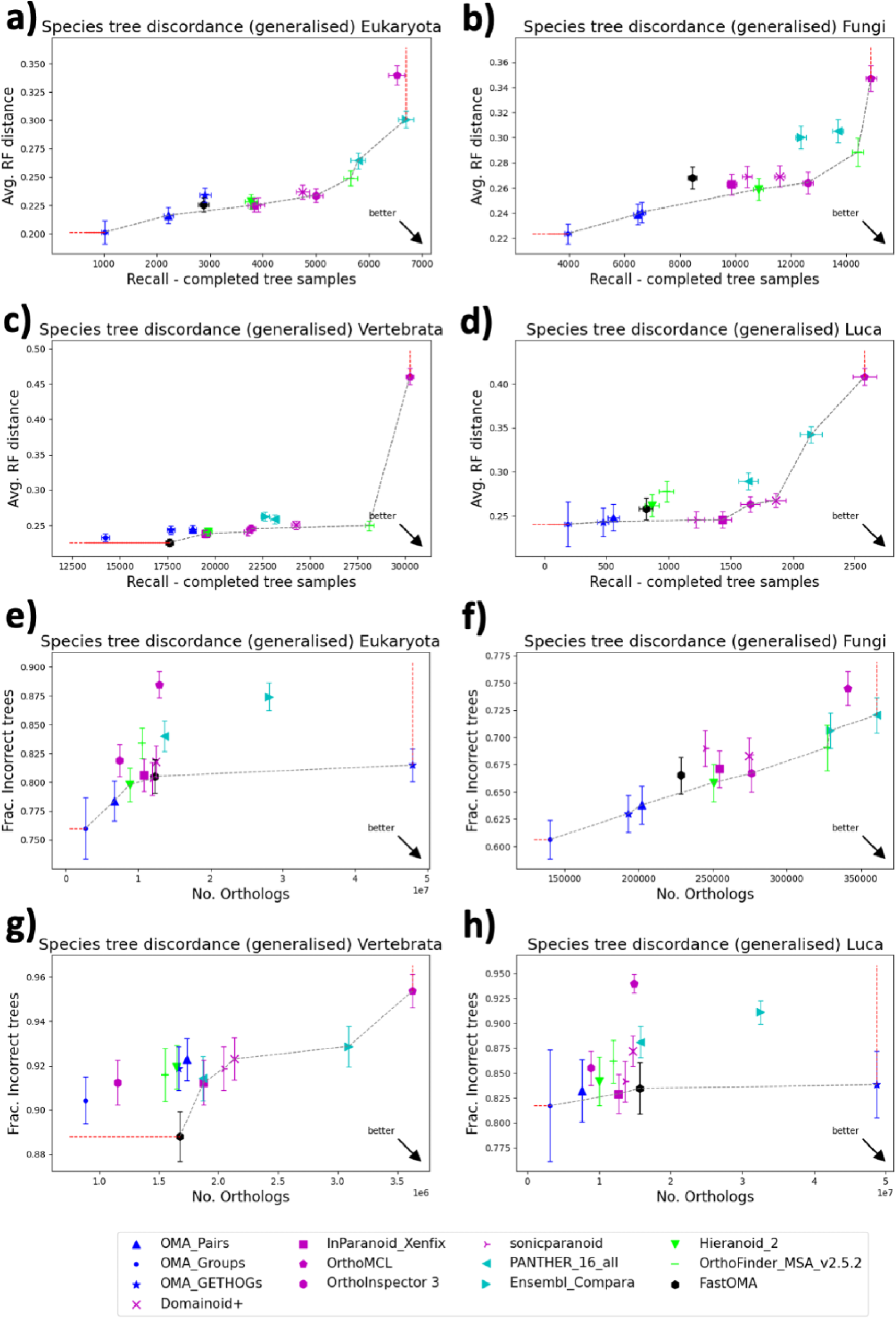
The result of generalised species tree discordance test in terms of average Robinson-Foulds distance vs number of completed tree samples (a-d) and in terms of the fraction of incorrect trees and number of orthologs (e-h). The other methods of OMA are in blue and the new **FastOMA** is in black. Graph-based methods (OrthoMCL, OrthoInspector, InParanoid, Sonicparanoid, and Domainoid+) are in purple and the tree-based methods are in cyan. The hybrid methods which use both gene tree and graph structure are in green.

**Supplementary Figure 5.**
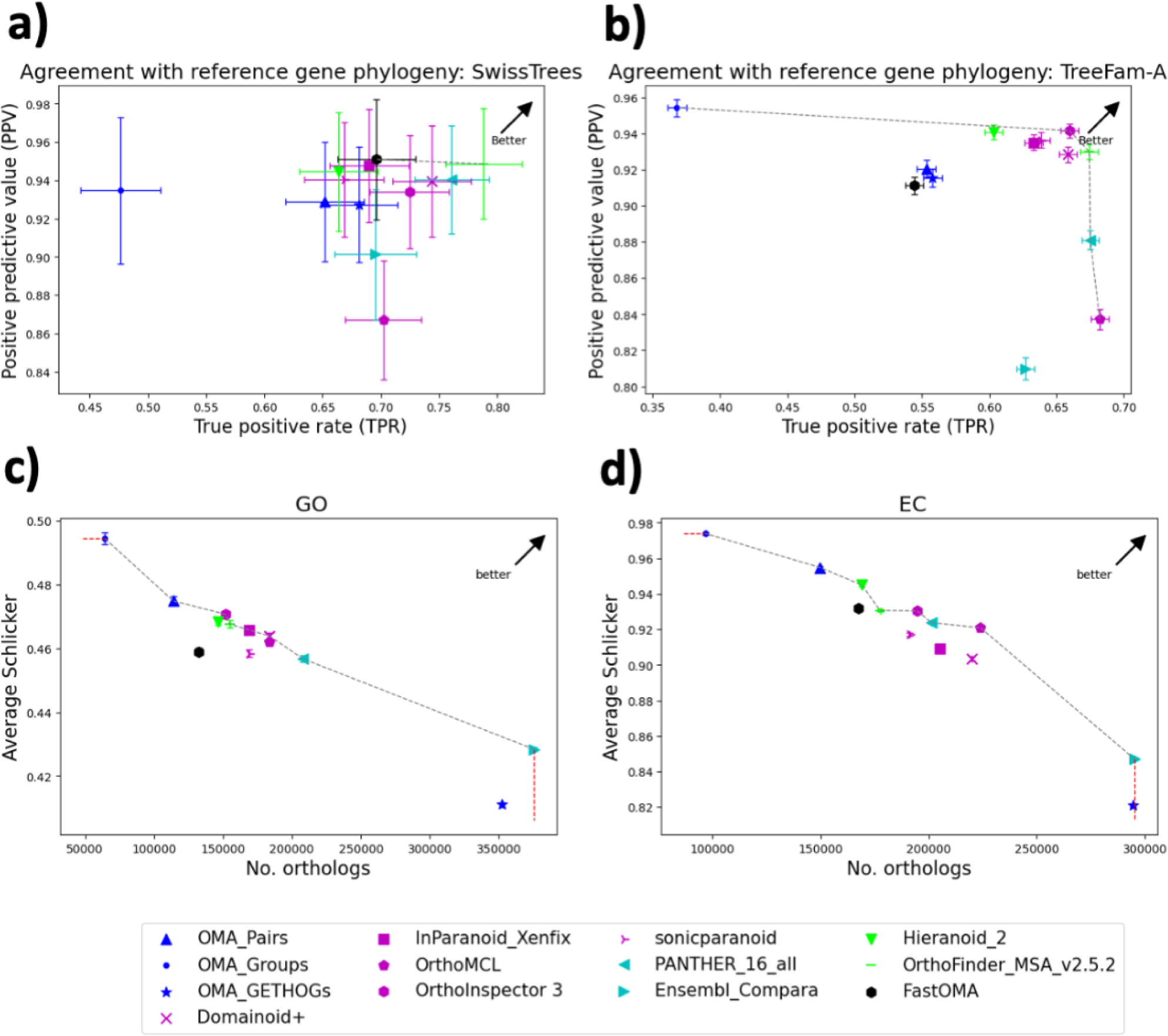
The result of agreement with reference gene phylogeny in terms of positive predictive value and true positive rate (a-b). The result of the Functional GO and EC tests in terms of average Schlicker (a similarity score) and number of orthologs (c-d).

## S3. Comparing FastOMA results with OMAmer mapping

FastOMA benefits from OMAmer mapping. However, mapping tools cannot provide enough information for orthology inference. To showcase the superiority of FastOMA against OMAmer mapping, we compared the results of QfO benchmarking tests. Note that the OMAmer tool is solely attributing genes to HOGs (gene family) and thus only predicts homology to other members of the gene family. It cannot differentiate paralogs from orthologs. To find the “orthologous” pairs using OMAmer mapping, we selected the gene with the highest OMAmer score of each species, for each HOG, and we generated orthologous pairs between genes from different species when they are attributed to the same HOG. Gene pairs from the same species are excluded since they are paralogous. We also generated orthologous pairs of genes where one is from a HOG and the other is from its parent HOGs. The results of the QfO species tree discordance benchmarking for such mapping alone show poor performance, with both lower recall and precision than FastOMA and most other orthology methods (**Supplementary Figures 6-8**). This shows the benefit of FastOMA’s post-OMAmer-mapping orthology inference algorithm.

## S4. Impact of reference HOG database on FastOMA results

Some of the QfO proteomes are already included in the OMAmer database of reference HOGs, which could introduce a bias in FastOMA’s favour. To study this effect, we removed such proteomes from the OMAmer database on which we run FastOMA, and used the species discordance benchmark to measure the extent of the bias. Overall, using a database where those proteomes are not present does not significantly affect the results from FastOMA, with most of the difference between versions being within error bars. The QfO results are reported in **Supplementary Figures 6-8**.

**Supplementary Figure 6.**
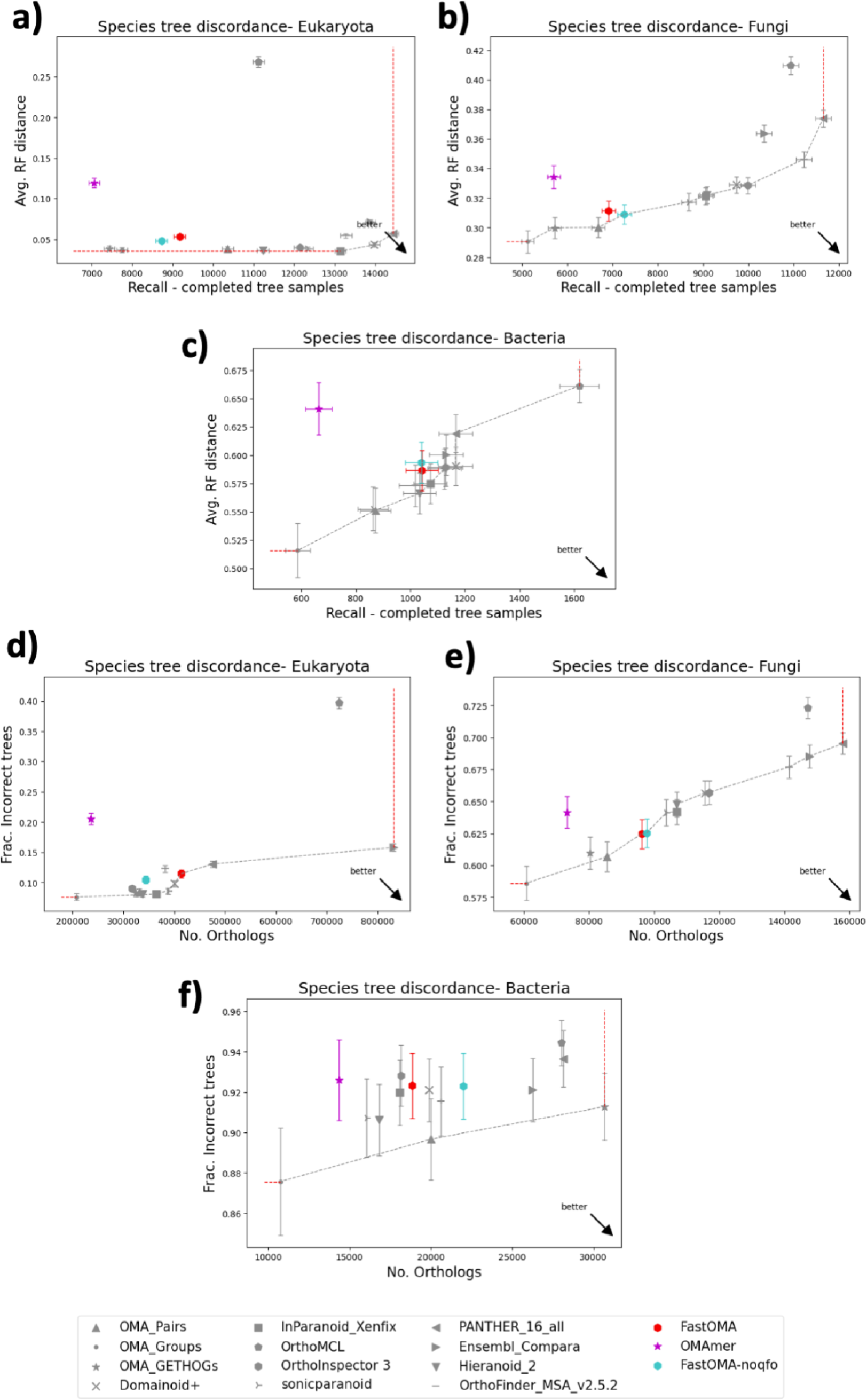
The result of the species tree discordance tests comparing OMAmer (pink) and FastOMA including (red)/excluding (cyan) QfO species in the reference set. (a-d): Benchmark results in terms of average Robinson-Foulds distance vs number of completed tree samples. (e-h): Benchmark results in terms of the fraction of incorrect trees and number of orthologs.

**Supplementary Figure 7.**
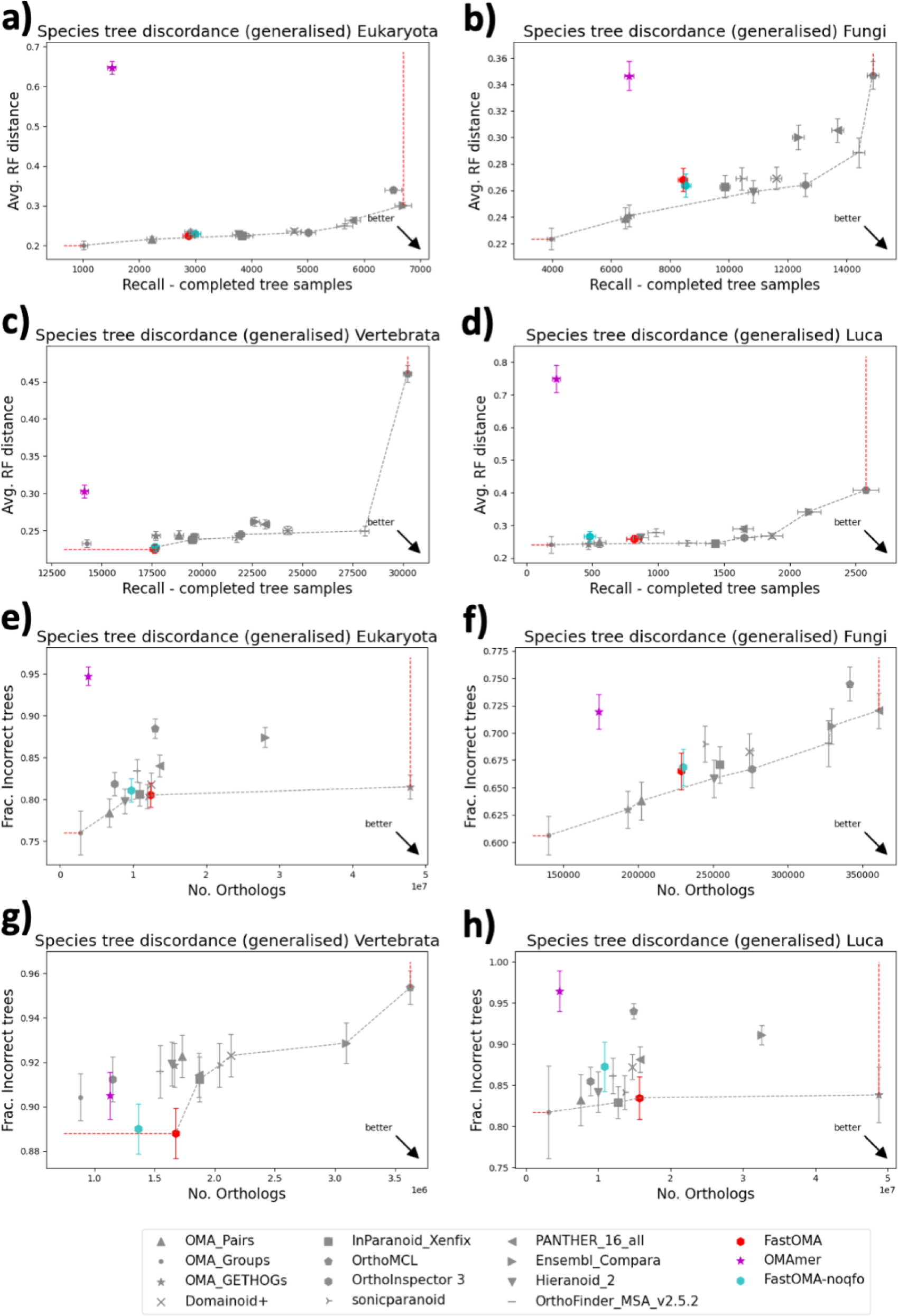
The result of the generalised species tree discordance tests comparing OMAmer (pink) and FastOMA including (red)/excluding (cyan) QfO species in the reference set. (a-d): Benchmark results in terms of average Robinson-Foulds distance vs number of completed tree samples. (e-h): Benchmark results in terms of the fraction of incorrect trees and number of orthologs.

**Supplementary Figure 8.**
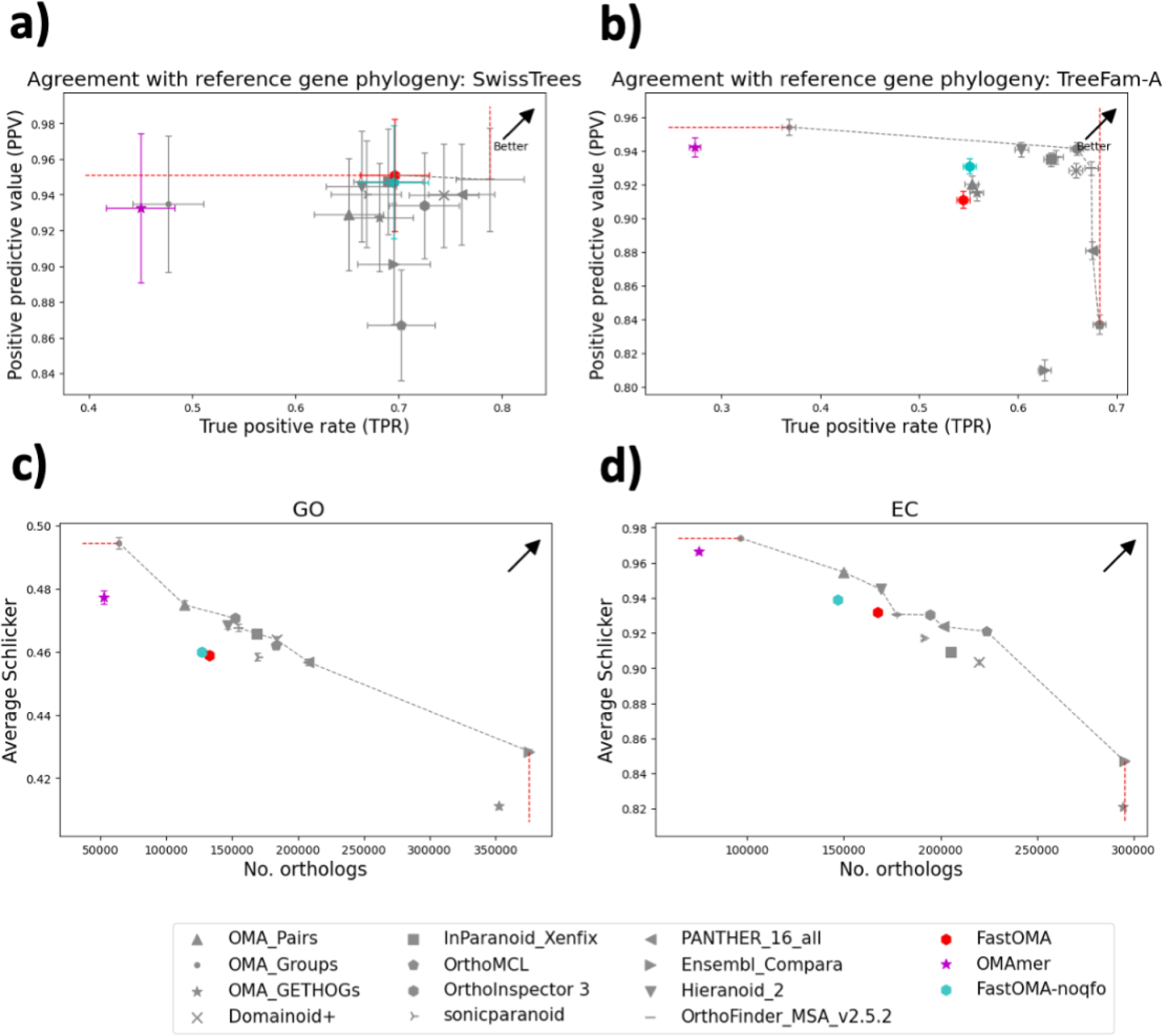
The result of the agreement with reference gene phylogeny tree tests and functional tests comparing OMAmer (pink) and FastOMA including (red)/excluding (cyan) QfO species in the reference set.

## S5. FastOMA robustness on threshold

To study the impact of threshold parameters used in FastOMA, we used different parameters and evaluated the results with the QfO species tree discordance tests. Specifically, we changed the thresholds of MSA trimming (row- and column-wise), the score of species overlap method for detecting duplication events and number of proteins sampled per subHOG. Results are reported in **Supplementary Figures 9-11** and **12-17**. The threshold changes only had a minimal effect on the results, with all results being between error bars. FastOMA with default parameters in most cases provide a better tradeoff in terms of precision and recall compared to the other tested parameters.

**Supplementary Figure 9.**
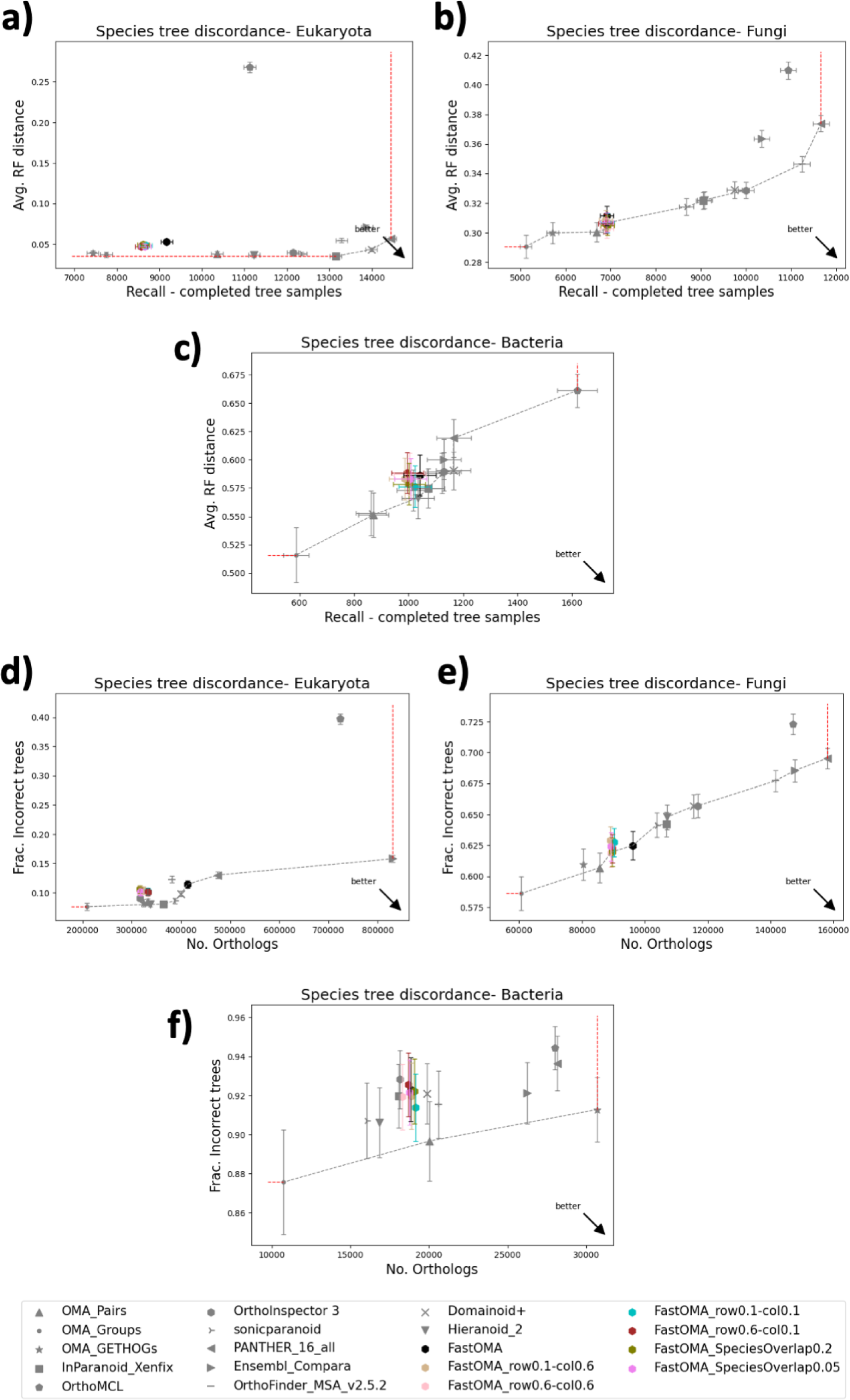
The result of species tree discordance tests considering different MSA trimming for rows and columns (with threshold of either 0.1 or 0.6) and species overlap scores of 0.2 or 0.05. FastOMA with default values is shown in black; MSA rows trimmed at 0.5, columns trimmed at 0.3, and a species overlap of 0.1.

**Supplementary Figure 10.**
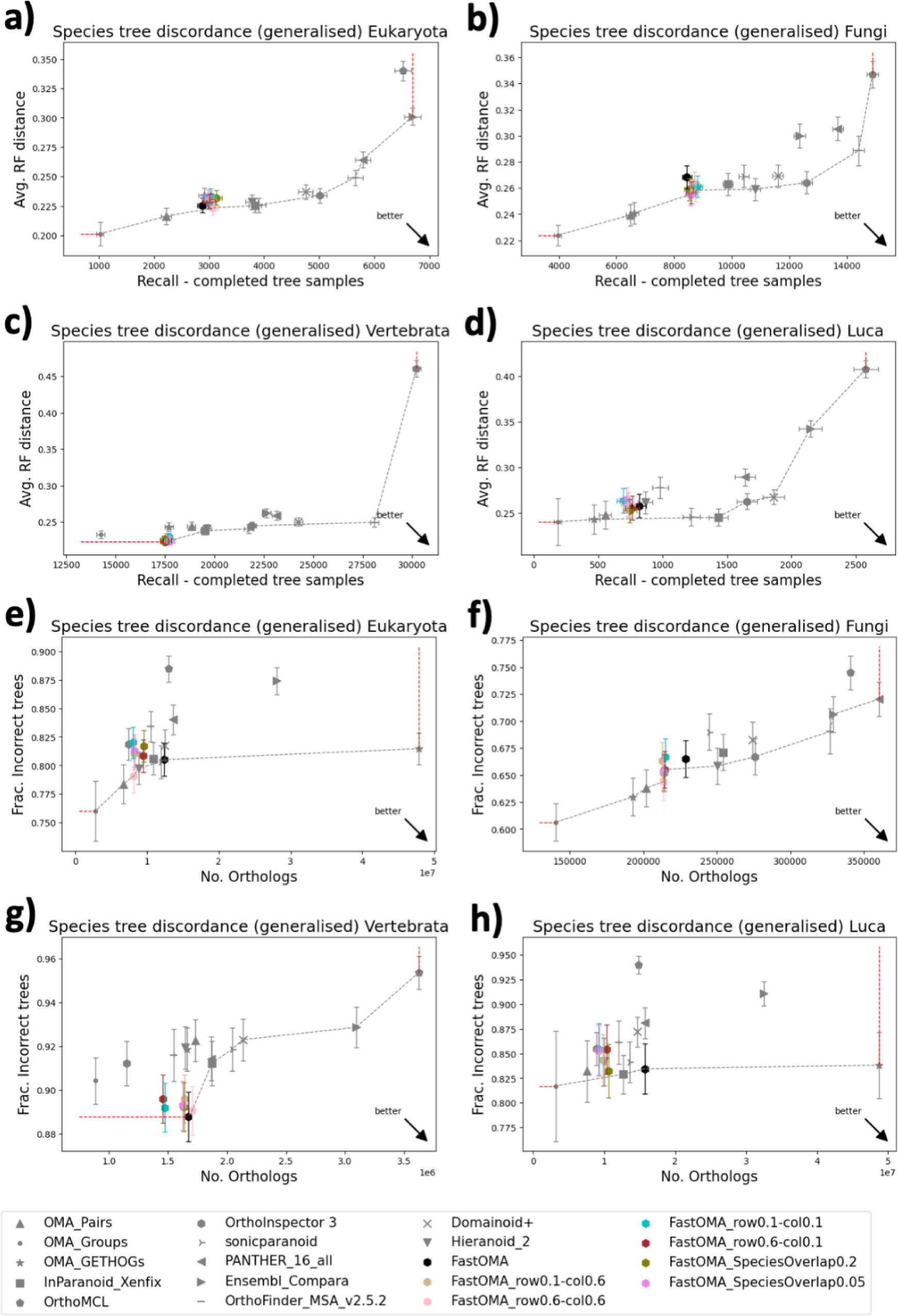
The result of generalised species tree discordance tests considering different MSA trimming for rows and columns (with threshold of either 0.1 or 0.6) and species overlap scores of 0.2 or 0.05. FastOMA with default values is shown in black; MSA rows trimmed at 0.5, columns trimmed at 0.3, and a species overlap of 0.1.

**Supplementary Figure 11.**
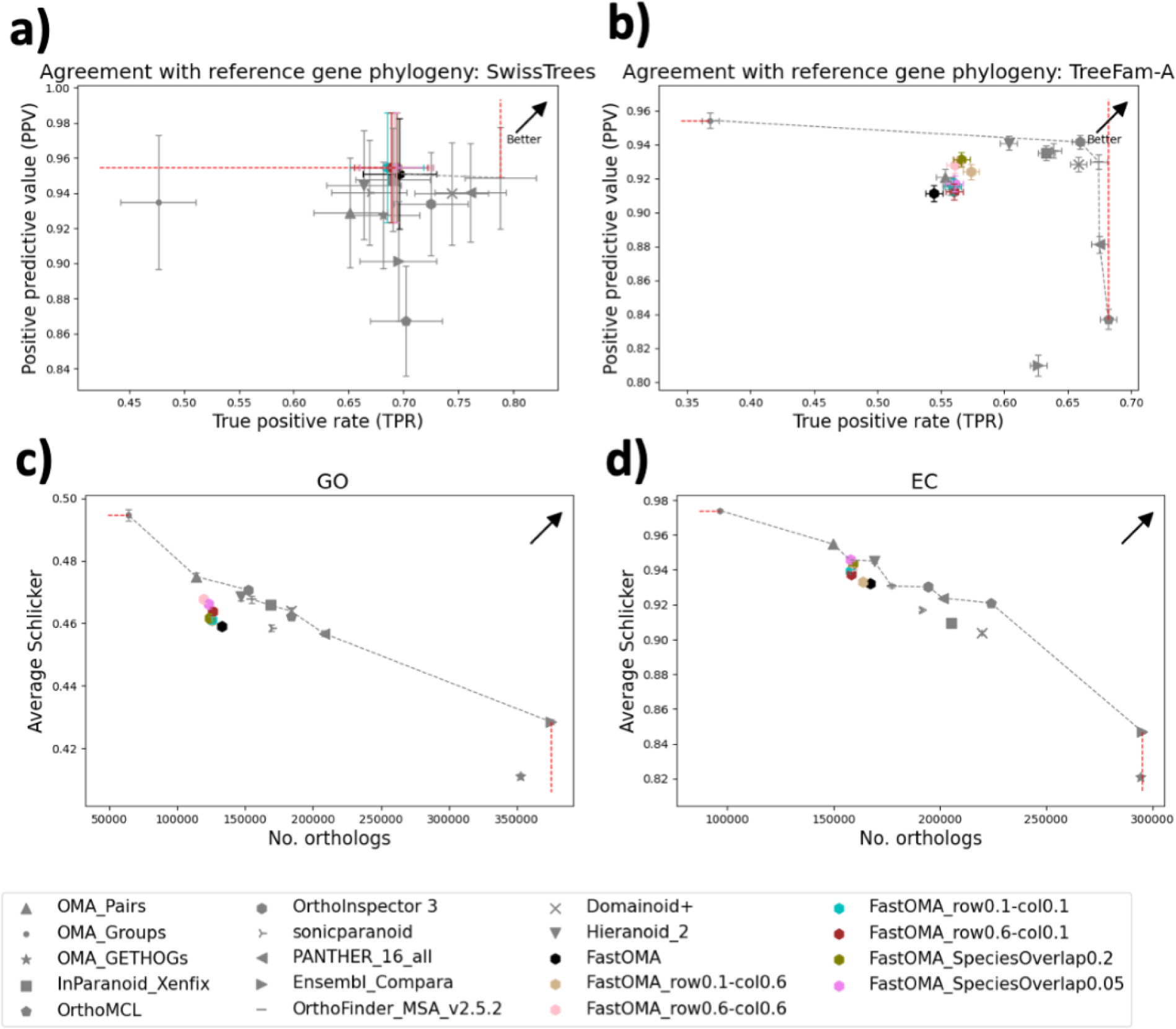
The result of agreement with reference gene phylogeny tree tests and functional tests considering different MSA trimming for rows and columns (with threshold of either 0.1 or 0.6) and species overlap scores of 0.2 or 0.05. FastOMA with default values is shown in black; MSA rows trimmed at 0.5, columns trimmed at 0.3, and a species overlap of 0.1.

**Supplementary Figure 12.**
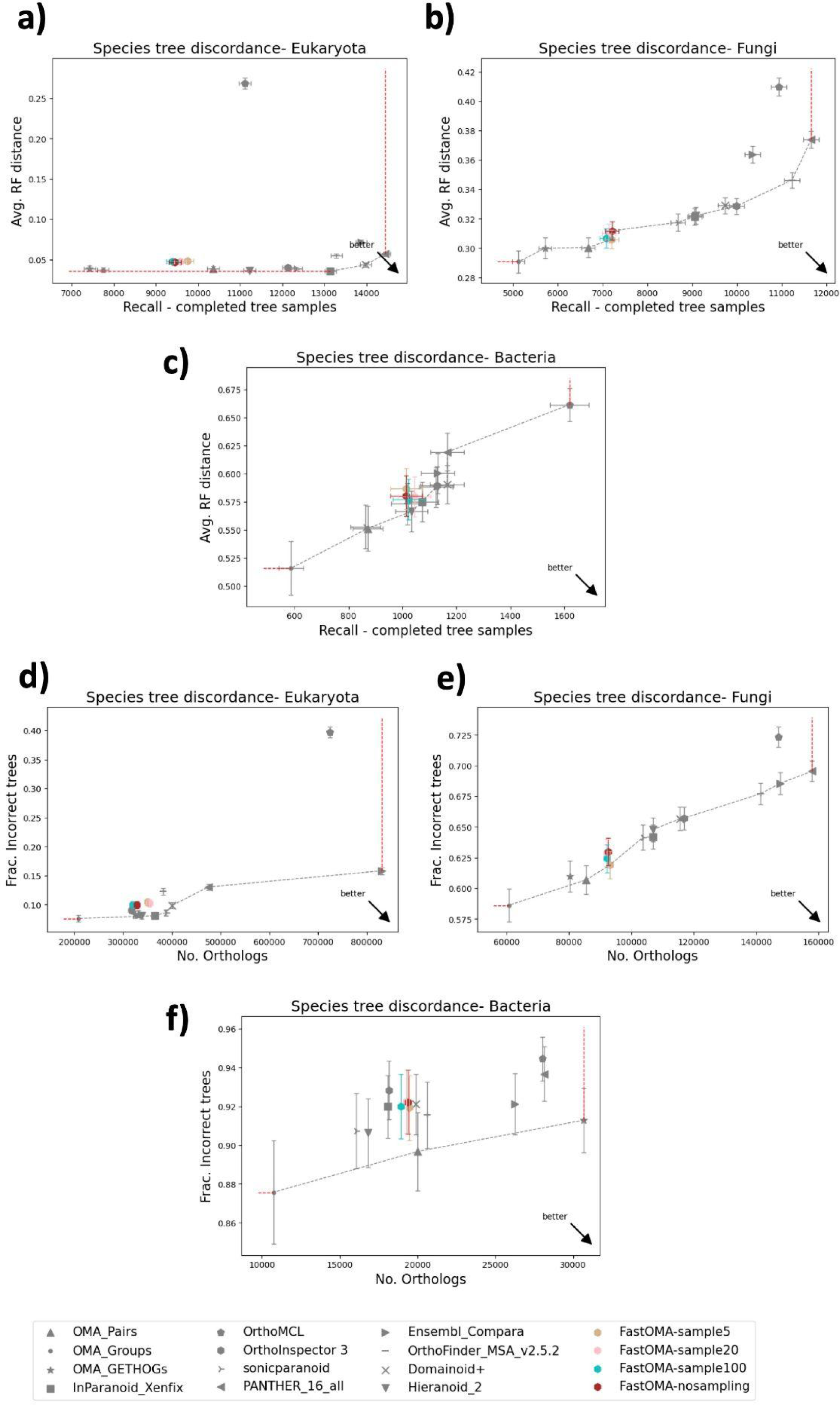
The result of species tree discordance tests considering different numbers of proteins used in subsampling 5, 20, and 100 and without subsampling.

**Supplementary Figure 13.**
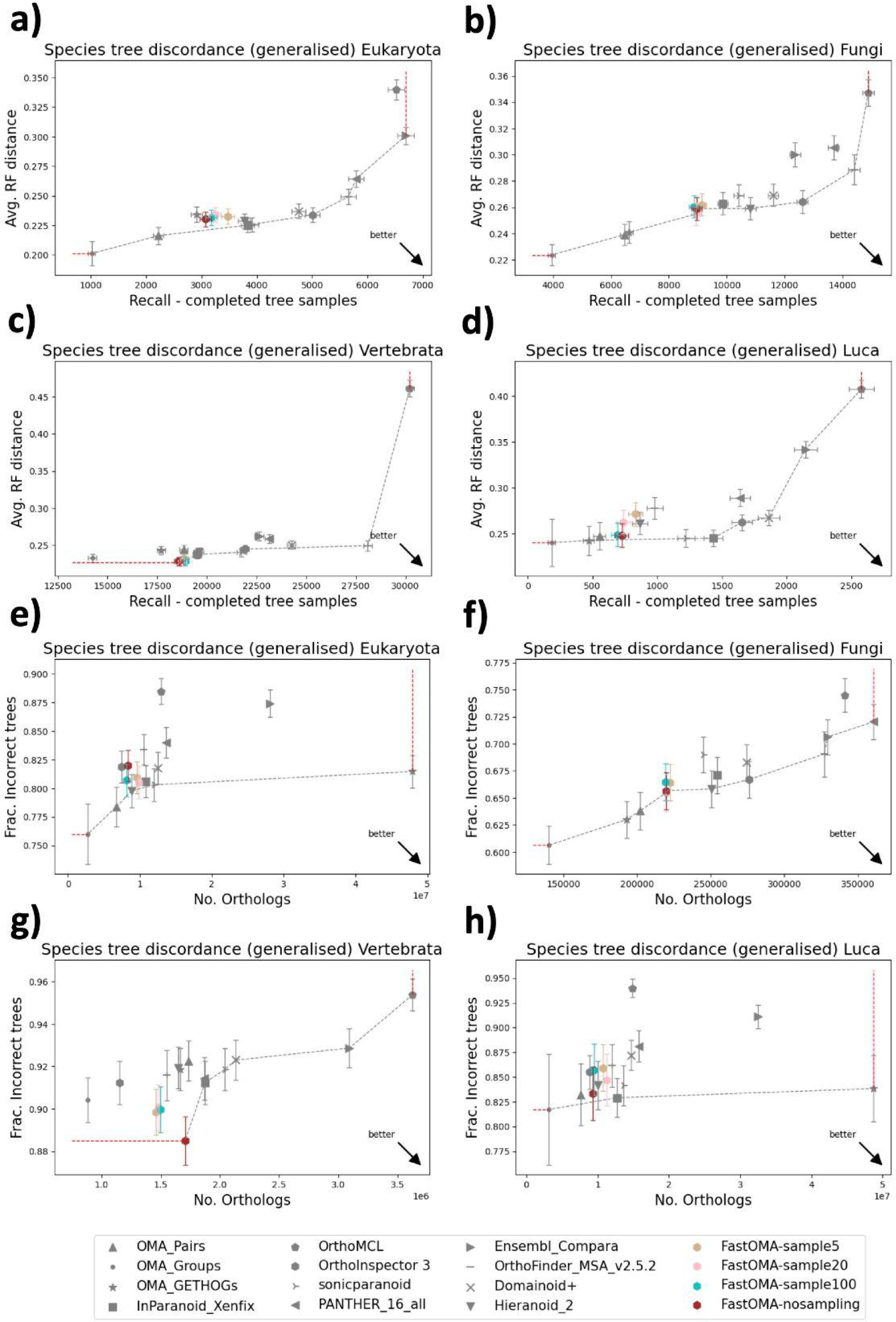
The result of generalised species tree discordance tests considering different numbers of proteins used in subsampling 5, 20, and 100 and without subsampling.

**Supplementary Figure 14.**
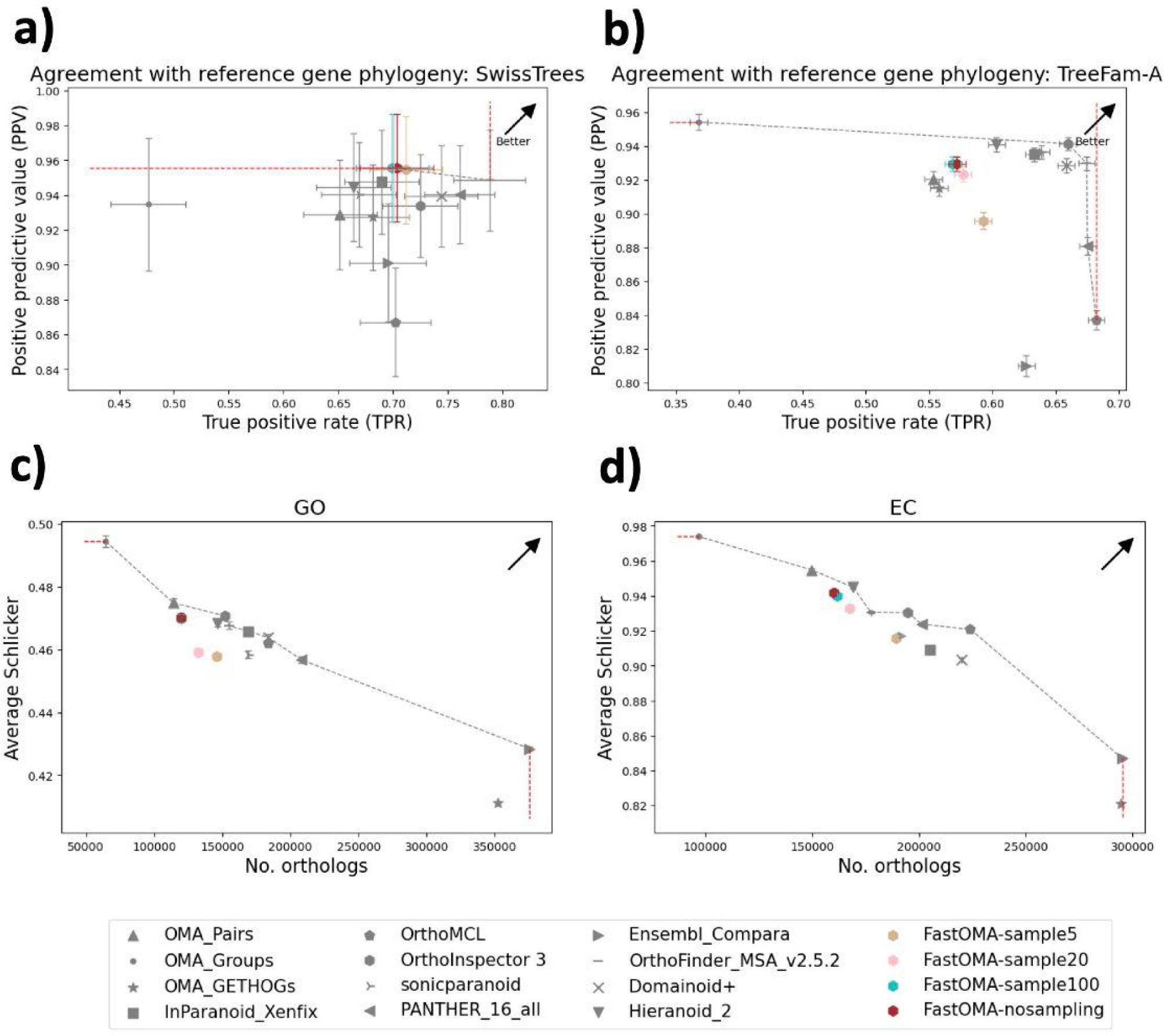
The result of agreement with reference gene phylogeny tree tests and functional tests considering different numbers of proteins used in subsampling 5, 20, and 100 and without subsampling.

## S6. Impact of OMA database on FastOMA

In this section, we describe the analysis to investigate how FastOMA is impacted by the reference OMA database. This is implemented by replacing the first step of FastOMA (i.e., OMAmer, which uses the OMA database) with the InterProScan package. The protein grouping (i.e., finding gene families) is a crucial step because if two orthologous genes are mapped to two different groups, there is no chance of rescuing these pairs in subsequent FastOMA steps. This results in false negatives, lowering the recall in orthology inference. Note that FastOMA’s grouping is a crucial step for achieving the speed; FastOMA only compares proteins that are inside the gene family, in contrast to other methods that do all-vs-all comparisons.

The benchmarking results provided in Supplementary Figures 15-17 show higher RF distance values and fraction of incorrect trees in most of the discordance tests when comparing FastOMA (OMA database) and FastOMA (InterProScan). However, FastOMA (InterProScan) was able to report more orthologous pairs and better recall. In reference gene families benchmarks, such higher recall was achieved at the expense of a drop in positive predictive values. Over all benchmarks, this strategy leads to overall higher recall but lower accuracy.

**Supplementary Figure 15.**
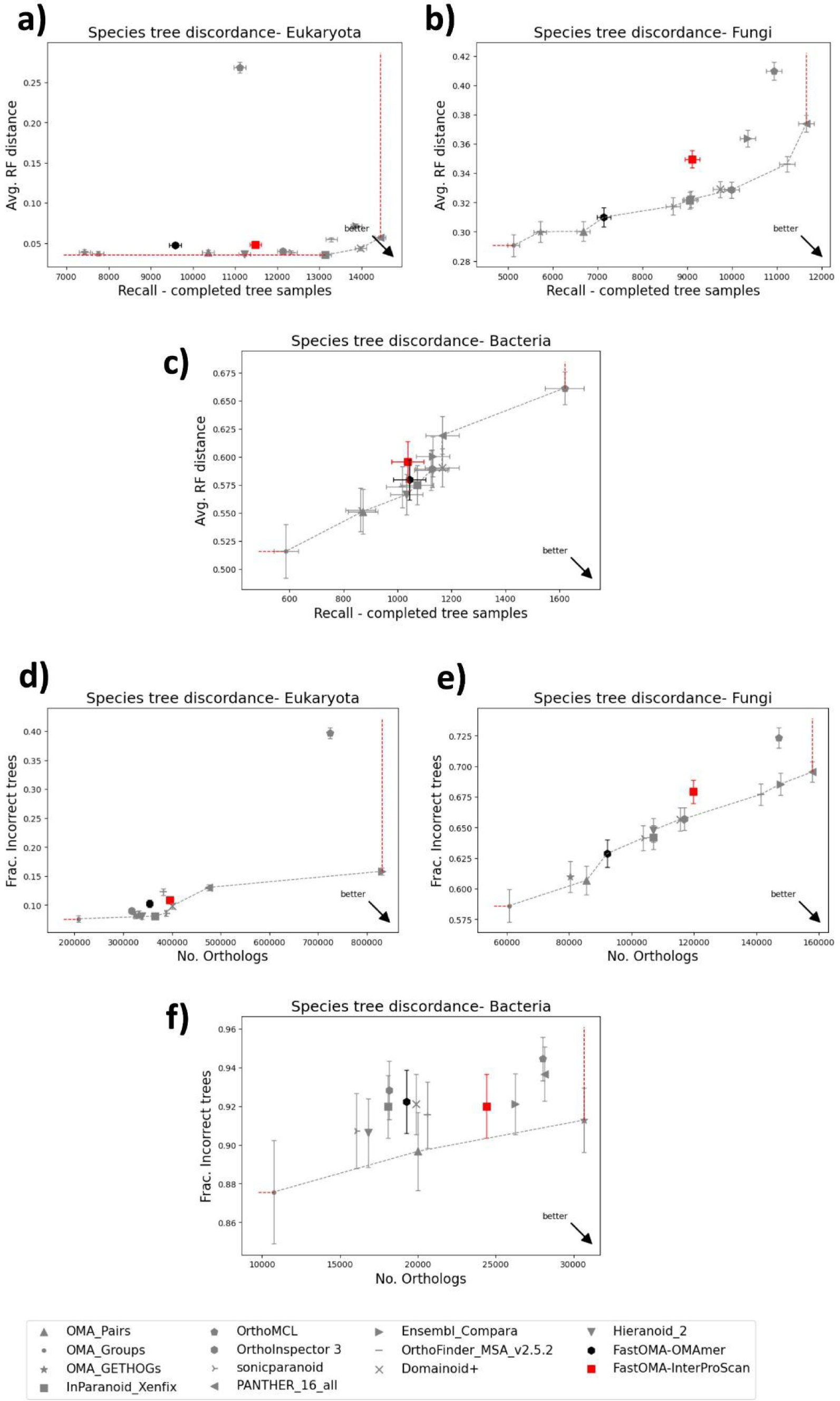
The result of species tree discordance tests running FastOMA on the InterProScan gene families (without the OMA database).

**Supplementary Figure 16.**
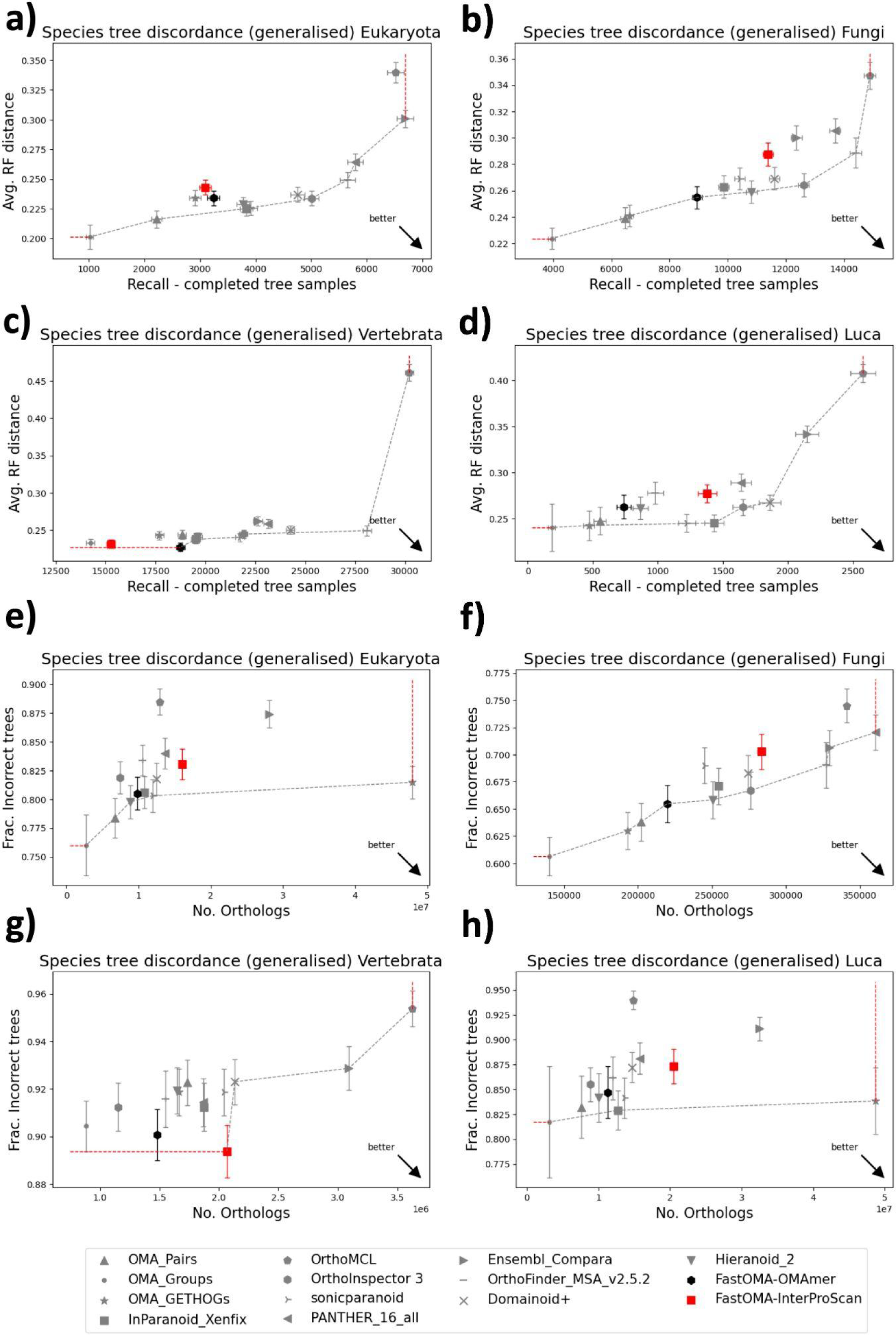
The result of generalised species tree discordance tests with FastOMA on the InterProScan gene families (without the OMA database).

**Supplementary Figure 17.**
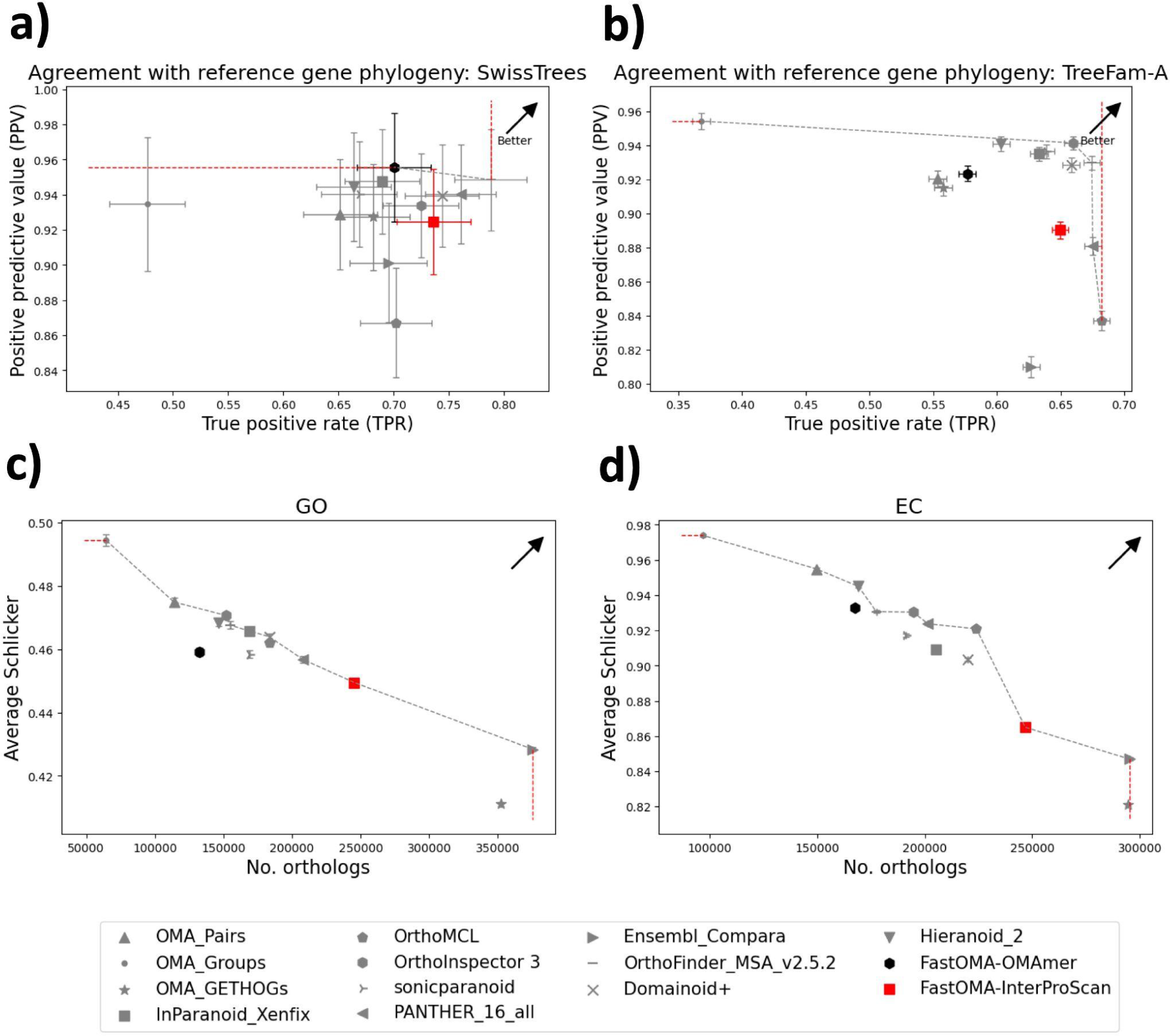
The result of agreement with reference gene phylogeny tree tests and functional tests with FastOMA on the InterProScan gene families (without OMA database).

## S7. Impact of species tree on FastOMA

Since one of the FastOMA’s inputs is the species tree, we investigated its impact on the orthology inference. As described in the Method section, we swapped five pairs of species (10 species in total) in the QfO dataset. The normalised Robinson-Foulds (RF) distance between the resulting trees and the true QfO species tree were 0.49 and 0.33 for “SwappedTree1” and “SwappedTree2”, respectively. This means that nearly half of the branches in SwappedTree1 were spurious. The results of benchmarking tests are shown in Supplementary Figure 18-20. Overall, this major perturbation of the input species tree did result in some performance deterioration, but to a relatively limited extent, and with the tree with more disruption (SwappedTree1) logically resulting in a worse outcome.

**Supplementary Figure 18.**
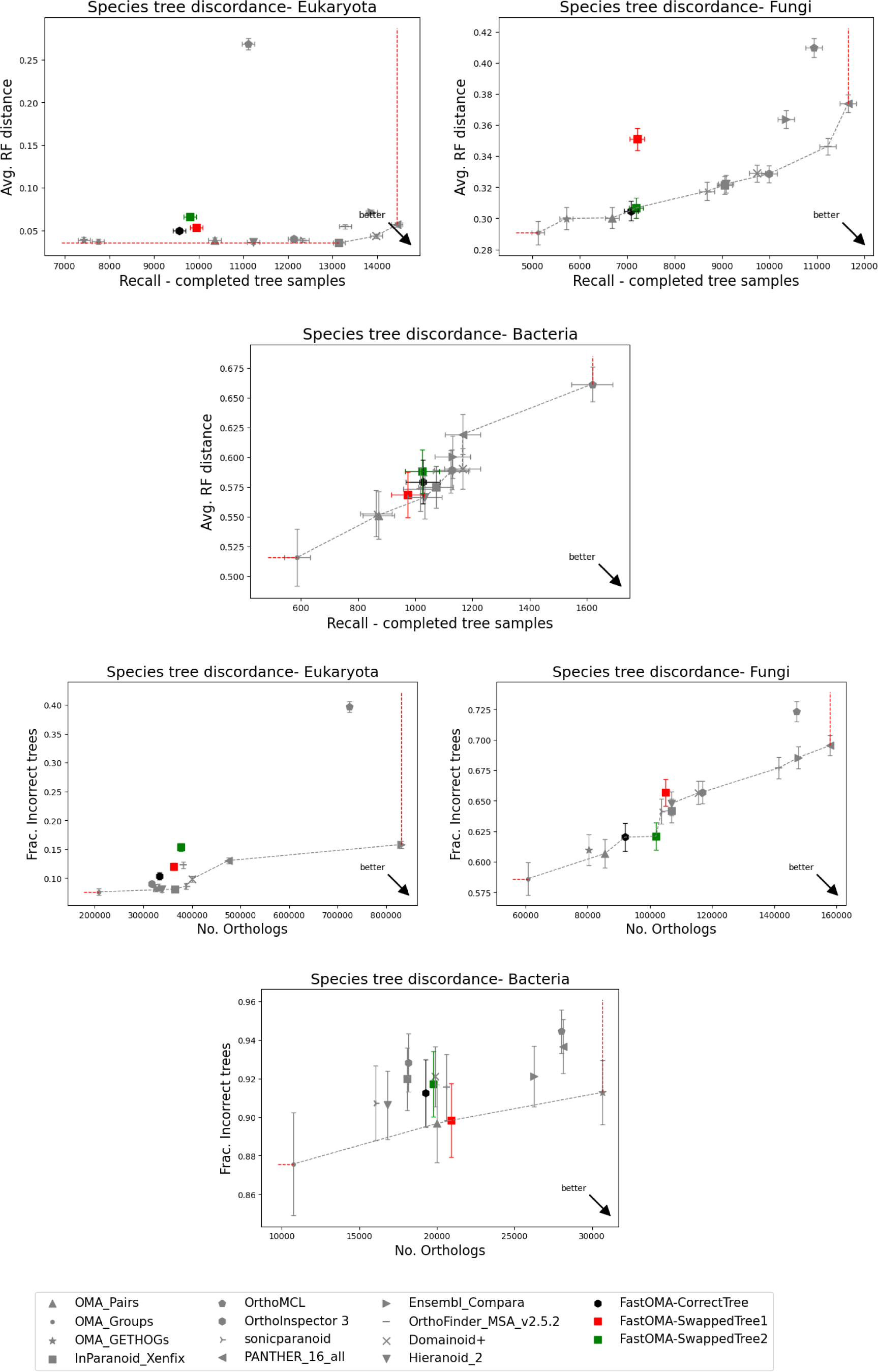
The result of species tree discordance tests running FastOMA with wrong species trees where five pairs of species are swapped.

**Supplementary Figure 19.**
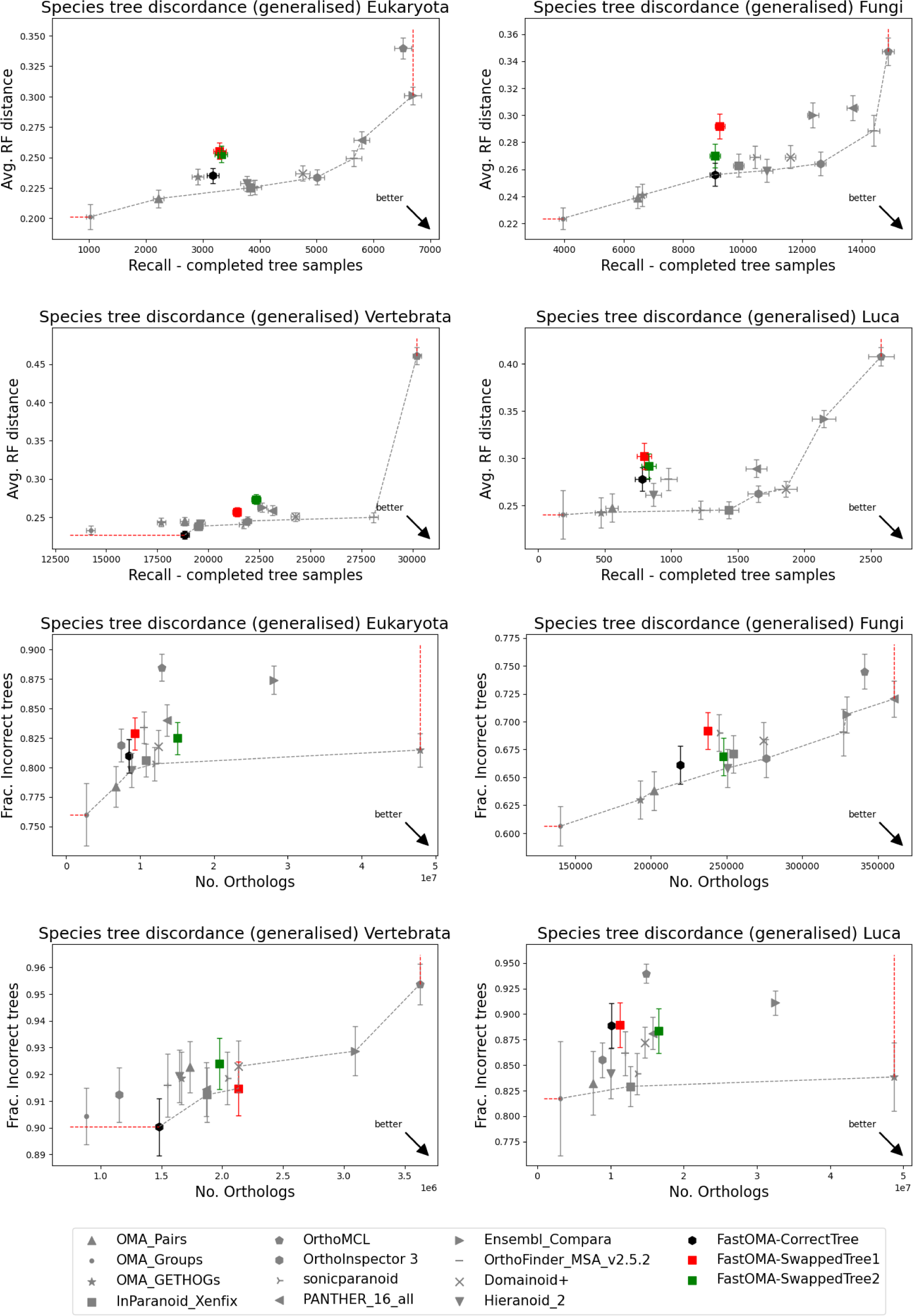
The result of generalised species tree discordance tests with wrong species trees where five pairs of species are swapped.

**Supplementary Figure 20.**
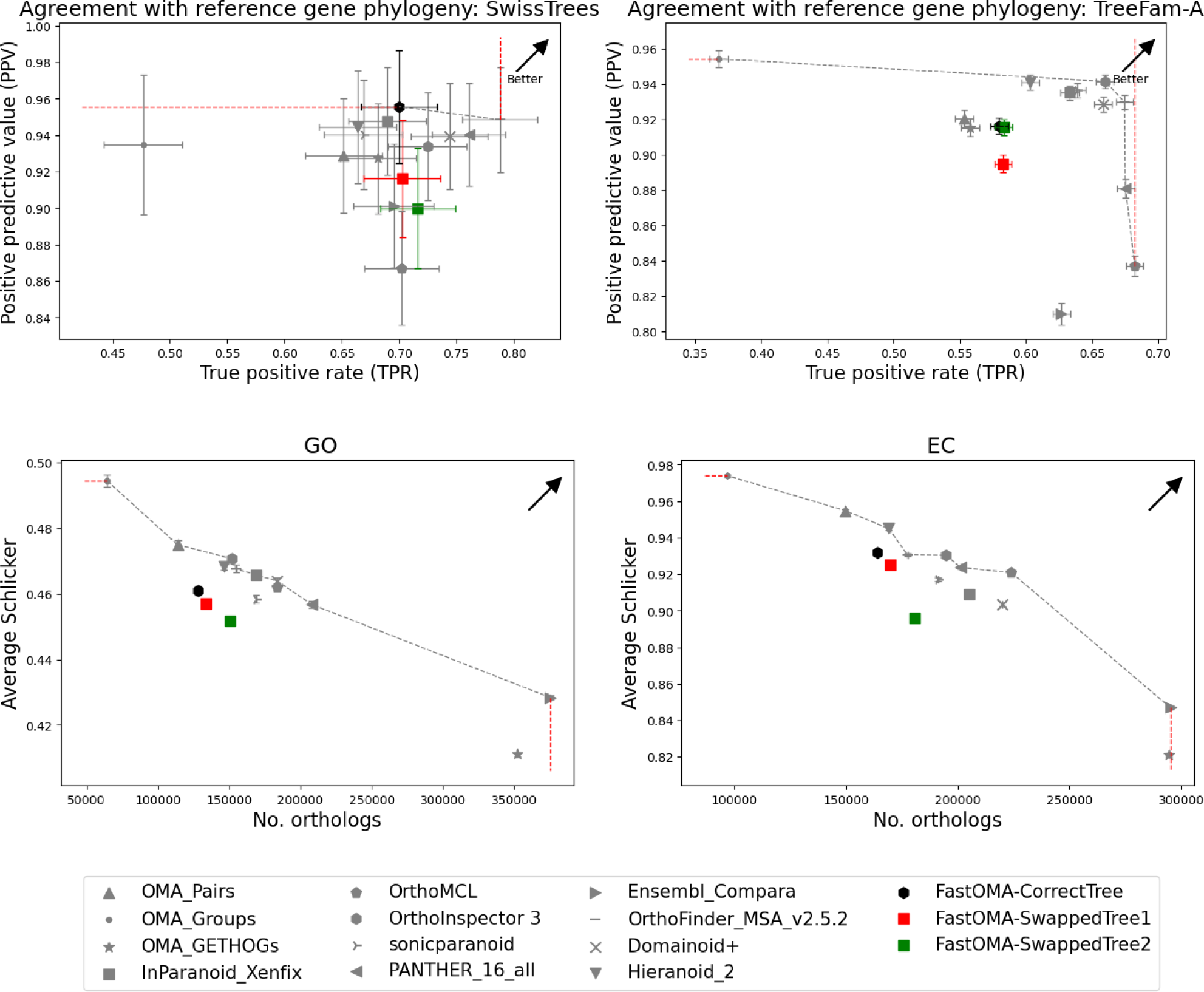
The result of agreement with reference gene phylogeny tree tests and functional tests with wrong species trees where five pairs of species are swapped.

## S8. The group benchmarking for the clade Bilateria

We also used the revisited Orthobench for benchmarking of orthologous groups^44^, which has been adapted as part of the QfO benchmarks. This benchmark assesses the ability of orthology inference to accurately predict 70 curated orthologous groups at the Bilateria level. As many of the tools in the QfO benchmark only report the orthology pairs, we could only include Panther and OMA-GETHOGs2 for comparison with FastOMA since these were the only two available as groups of proteins on the QfO public repository. FastOMA has a precision of 0.758 and 0.46 recall; this is a lower precision than OMA-GETHOGs2 with a slightly higher recall, but a higher precision than Panther which has a higher recall.

**Supplementary Table 1.** The result of group-based benchmarking for the clade Bilateria.

|  | Panther | OMA-GETHOGs2 | FastOMA |
| --- | --- | --- | --- |
| PPV (Positive predictive value, precision) | 0.58 | 0.876 | 0.802 |
| TPR (True positive rate, recall, sensitivity) | 0.56 | 0.43 | 0.518 |

## S9. FastOMA’s ability to select isoforms

In contrast to most other orthology methods, FastOMA considers multiple input alternative splicing isoforms and aims to identify the evolutionarily best-conserved one for orthology inference. FastOMA selects the isoform with the highest OMAmer family score, i.e., the one with the best k-mer similarity with its closest gene family given its length. We compared the results of FastOMA using different ways to select isoforms: choosing the longest one as is often done by other methods, selecting the UniProt reference isoform, and FastOMA’s selection. The analysis on UniProt reference proteomes showed that FastOMA’s selection resulted in the most parsimonious results, i.e., the least number of rootHOGs and total implied losses (**Supplementary Figure 12**) when reconstructing gene family evolutionary histories. FastOMA selection resulted in the non-longest isoforms being selected for 35% of the proteins with multiple isoforms.

**Supplementary Figure 21.**
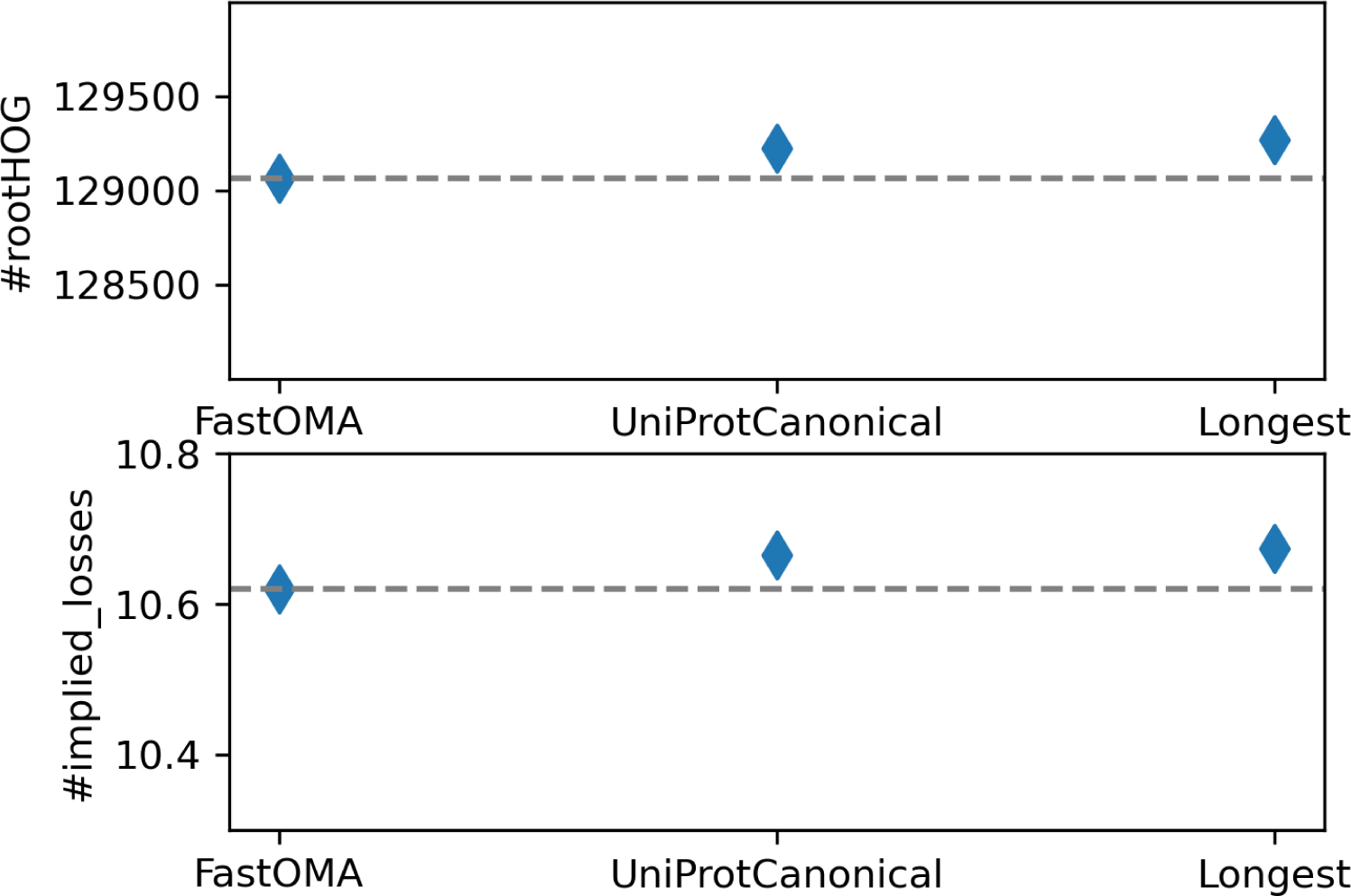
Impact of selecting the isoforms on the number of rootHOGs and the total number of implied losses comparing FastOMA’s selection with the UniProt Canonical and longest isoforms.

## S10. FastOMA’s ability to find split genes

FastOMA is capable of finding split genes, i.e., parts of the same gene predicted as multiple different genes, and merging them in the multiple sequence alignment (MSA). This is done to correct issues that might arise due to errors in genome annotation or to fragmented genome assembly where a gene is split across separate scaffolds; two complementary parts of the same gene being included in an MSA would result in an incorrect tree and incorrect labelling of speciation events. Split genes (gene fragments) are found by comparing pairs of genes (rows) in the MSA and those row pairs with complementary gaps and with an overlap in the MSA of less than 15% of alignment length are considered as candidates. These candidate pairs are reported as split genes if they are closer to each other on the gene tree than one fifth of the maximum distance between two leaves of the gene tree, to avoid merging fragments of distant paralogs. They are then merged and considered as a single sequence from the rest of the FastOMA inference, and are reported as such in FastOMA’s OrthoXML output. In the UniProt Eukaryote reference proteomes, FastOMA identified 40,297 pairs of sequences (out of 34.4 million sequences) that are likely fragments of split genes, most often found in species with a high proportion of fragments as detected by OMArk and BUSCO. Flagging these split genes aids in cleaning genomic datasets for orthology inference by using more reliable sequences, which in turn will result in a better understanding of genomic architecture and evolution (**Supplementary Figure 22**).

**Supplementary Figure 22.**
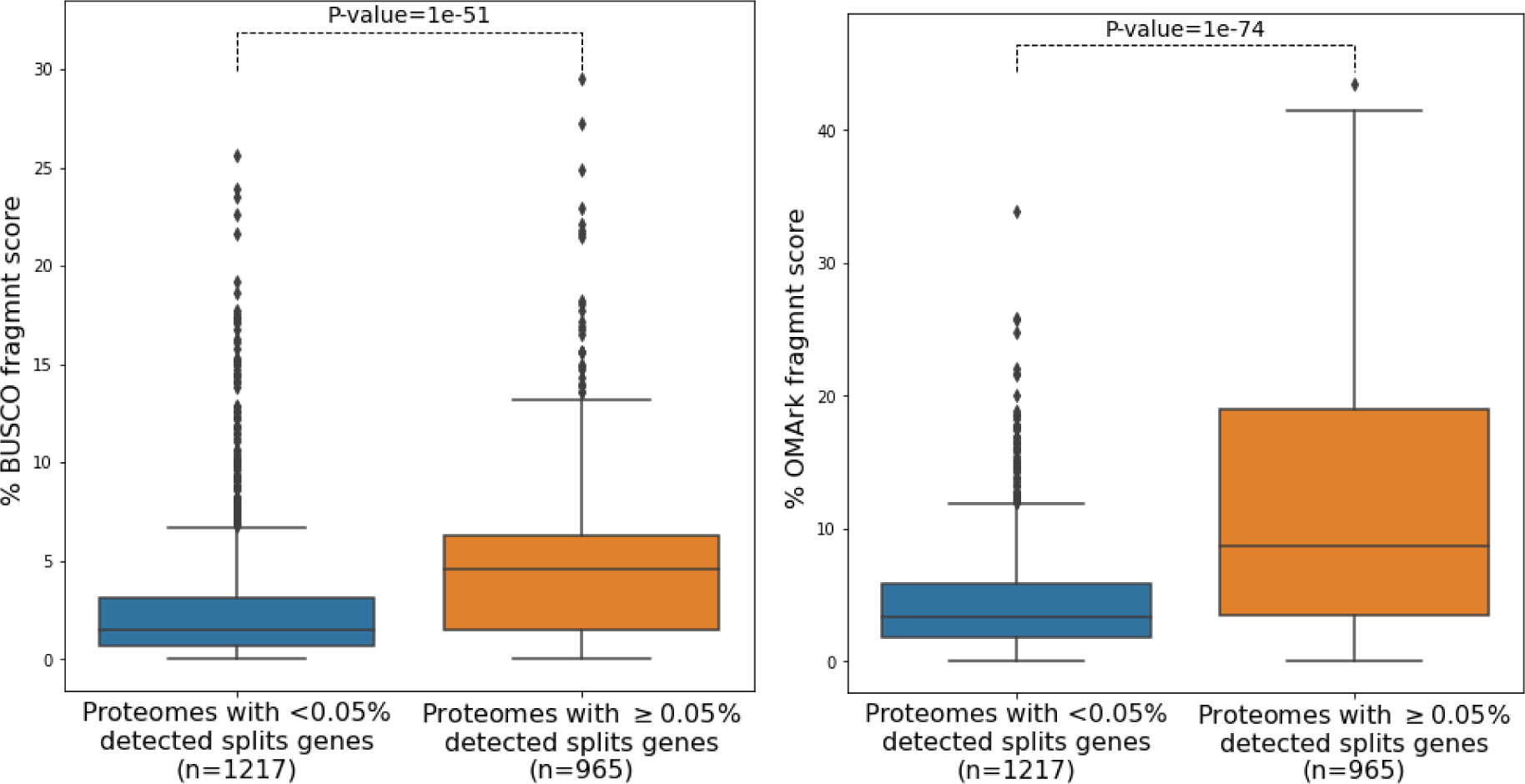
FastOMA’s ability to find split genes. The y-axis shows the percentage of fragments in proteomes estimated by BUSCO (left) and OMArk (right), partitioned into two groups of proteomes with higher or lower than 0.05% fragments found and merged by FastOMA. The result of the two-sided Mann-Whitney U rank test is reported on top of each figure.

## S11. FastOMA Nextflow DAG

FastOMA benefits from four sub-packages, written in Python. These include *check-input*, *infer-roothogs, batch-roothogs, infer-subhogs,* and *collect-subhogs.* These subpackages are used in a Nextflow pipeline and are dependent on each other through a directed acyclic graph (DAG), visualised below.

As an example, consider a dataset with 10 species and 1004 reference gene families. First, the job *check_input* with one CPU core is submitted. Once finished, 10 *OMAmer* jobs each with one CPU core will be run. *FastOMA.nf* waits until all of them are done. Then, using one CPU core, *infer_roothog* groups the proteins into 1004 families. Next, using one CPU core, *batch_rootHOG* generates (for example) 5 batches of small rootHOGs (each of which includes 200 rootHOGs) and 4 big rootHOGs. Now, *infer_subHOG (*under the wrapper *hog_rest)* starts for the 5 batches (each with one CPU core) and 4 big rootHOGs (under the wrapper *hog_big,* each with 6 CPU cores) totaling 5+24=29 CPU cores. Once all batches are finished, *collect_suhog* reads the output of each batch and writes the final OrthoXML file.

If the total number of needed CPU cores exceeds 500, Nextflow submits the first 500 jobs and sequentially adds one more job, once a job finishes. Start of a job relies on the availability of computation nodes assigned with Slurm Workload Manager on a High Performance Computing cluster. When a job finishes with an error due to time/memory limits, FastOMA re-submits it by doubling the CPU, time or memory. FastOMA also tries to decide the number of CPU cores and memory based on the size of the gene families in the batch. The number of CPUs per task can be changed with the Nextflow configuration file available at our GitHub *FastOMA/conf/base.config*.

In short, FastOMA tries to parallelise the steps as much as possible (with the goal of optimising the wall-clock time) considering the dependencies, using a different number of CPU cores for each step, specified based on the number of species and gene families.

**Supplementary Figure 23.**
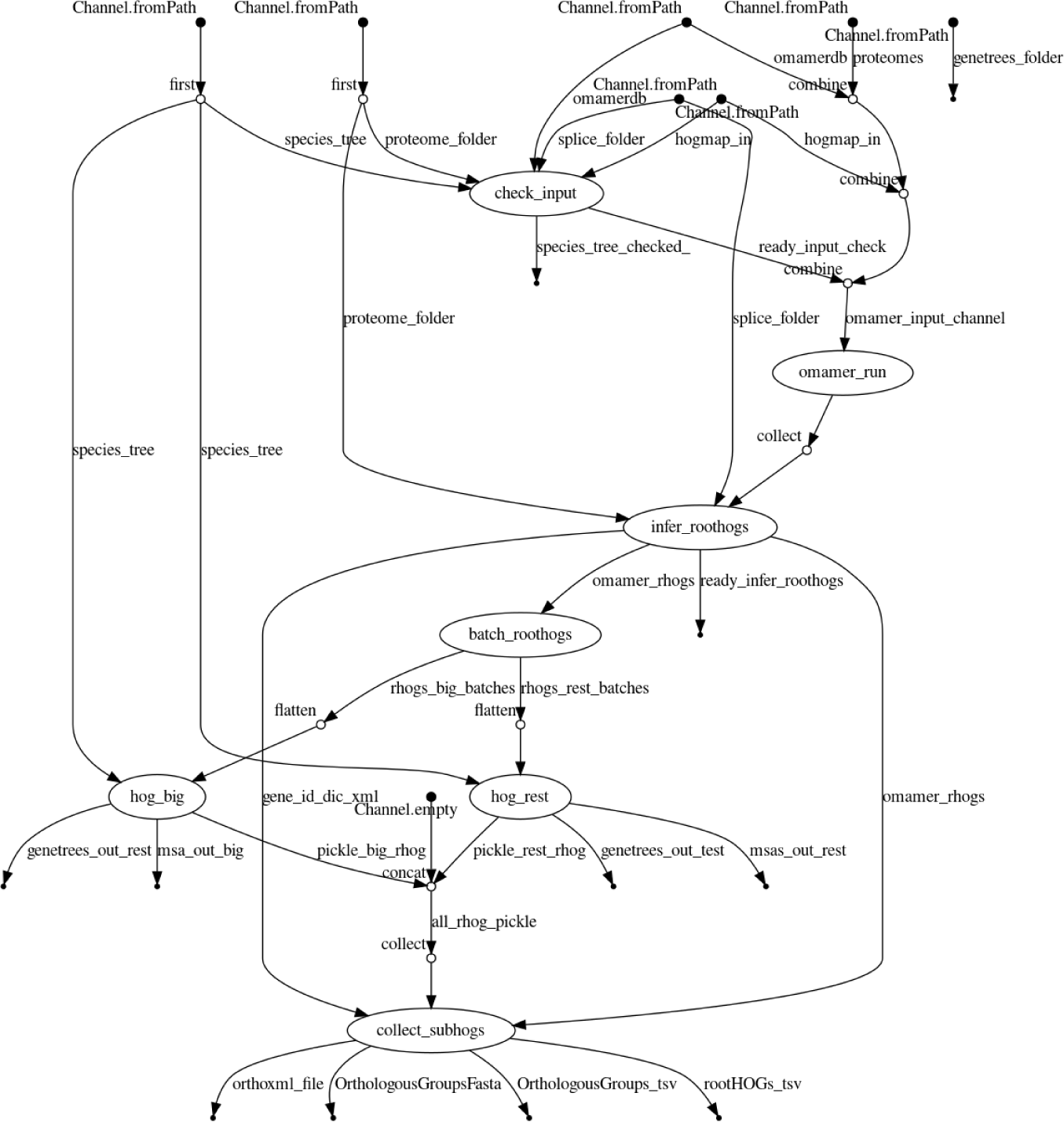
Subpackages of the FastOMA software.

**Supplementary Figure 24.**
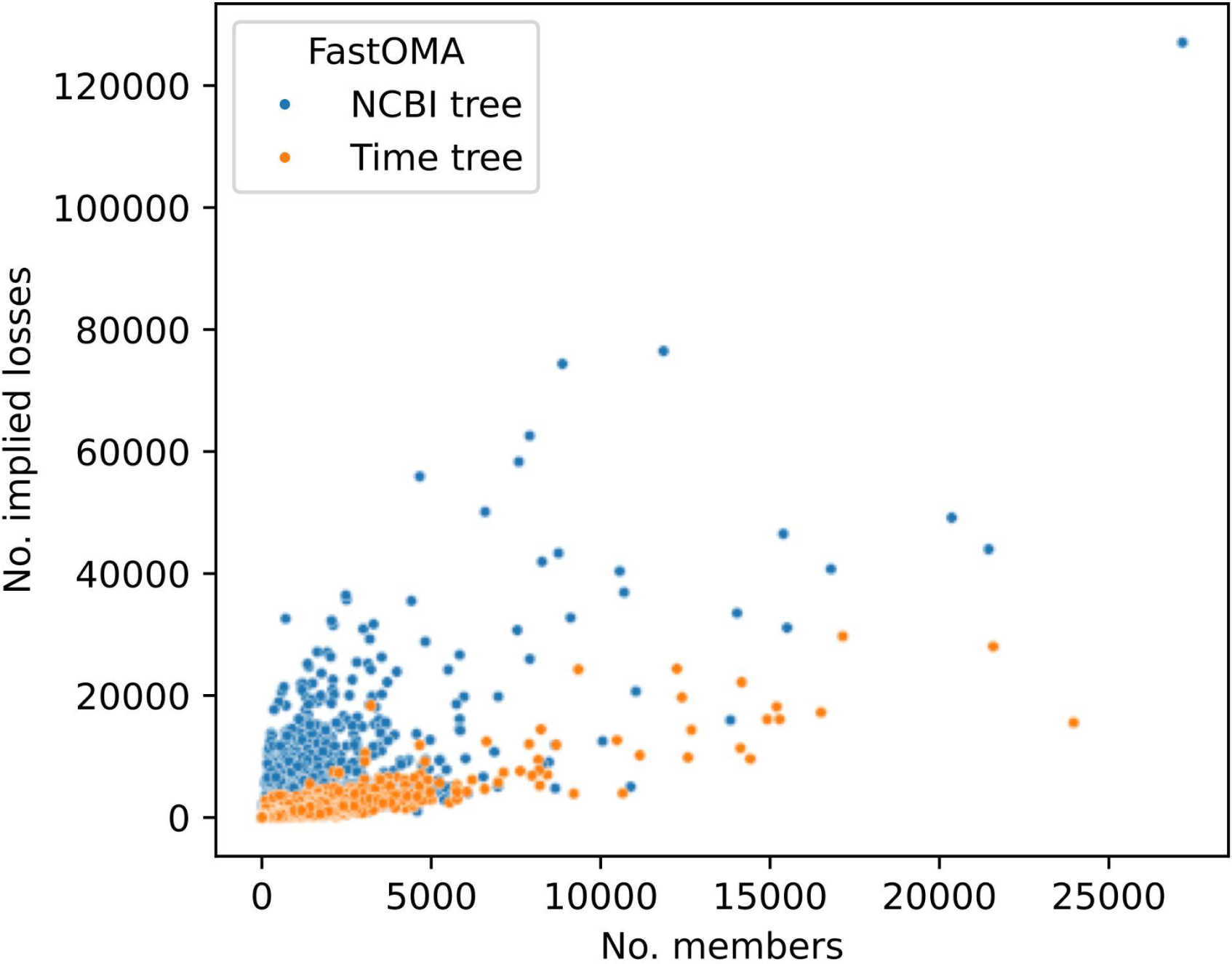
The impact of species tree resolution on evolutionary events in terms of implied losses. This is the full data of the figure shown in Figure 2d. Each point corresponds to a rootHOG. Number of rootHOGs that FastOMA found using the NCBI tree is 39,4516 and 38,5697 with the TimeTree.

**Supplementary Figure 25.**
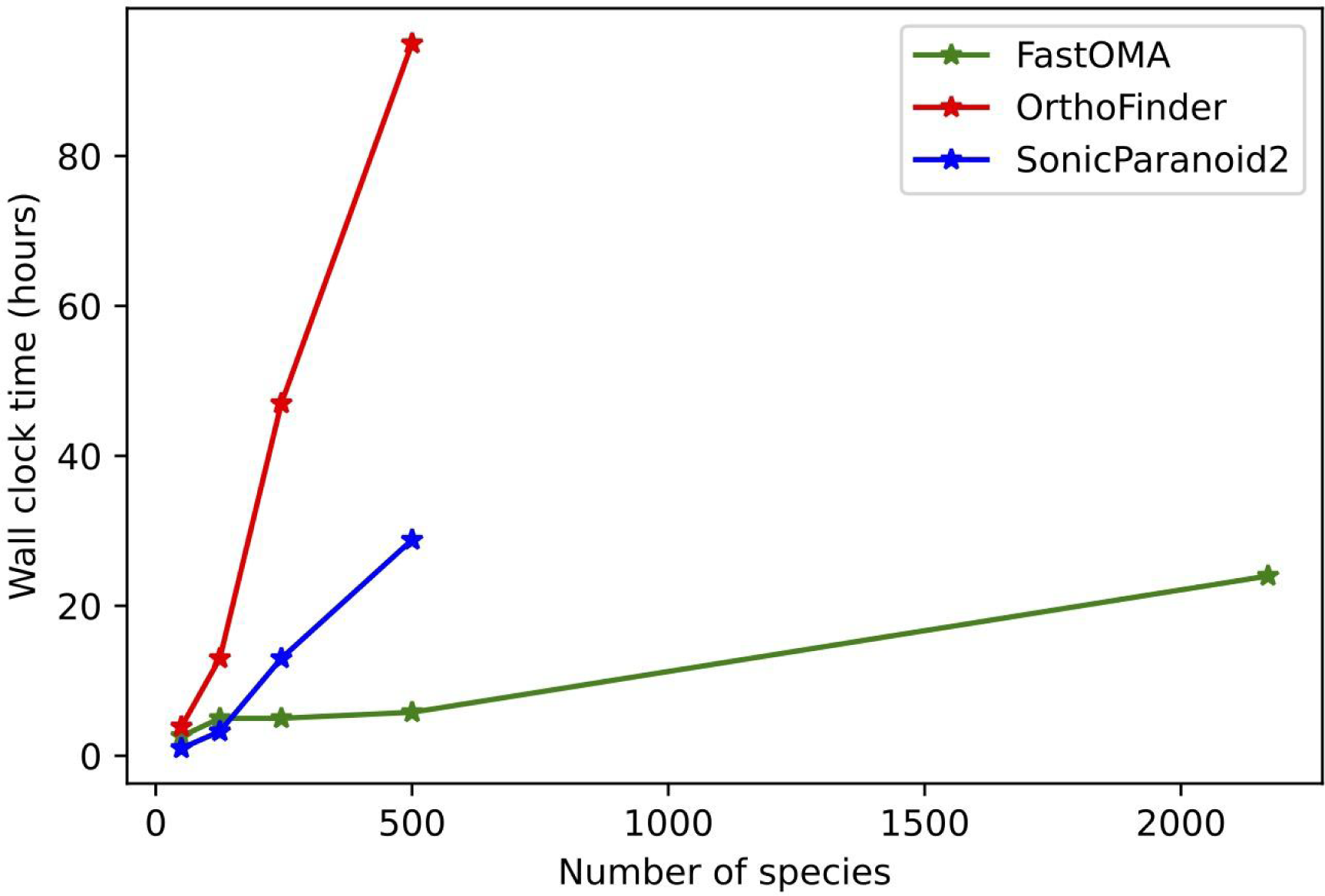
The comparison of tools in terms of Wall-clock time in hour.

## References

1. Lewin, H. A. et al. Earth BioGenome Project: Sequencing life for the future of life. Proc. Natl. Acad. Sci. U. S. A. 115, 4325–4333 (2018).

2. Fitch, W. M. Distinguishing homologous from analogous proteins. Syst. Zool. 19, 99–113 (1970).

3. Glover, N. et al. Advances and Applications in the Quest for Orthologs. Mol. Biol. Evol. 36, 2157–2164 (2019).

4. Linard, B. et al. Ten years of collaborative progress in the Quest for Orthologs. Mol. Biol. Evol. (2021) doi:10.1093/molbev/msab098.

5. Altenhoff, A. M. et al. OMA orthology in 2024: improved prokaryote coverage, ancestral and extant GO enrichment, a revamped synteny viewer and more in the OMA Ecosystem. Nucleic Acids Res. (2023) doi:10.1093/nar/gkad1020.

6. Dessimoz, C. et al. OMA, A Comprehensive, Automated Project for the Identification of Orthologs from Complete Genome Data: Introduction and First Achievements. in RECOMB 2005 Workshop on Comparative Genomics (eds. McLysaght, A. & Huson, D. H.) 61–72 (Springer-Verlag, 2005).

7. Kirilenko, B. M. et al. Integrating gene annotation with orthology inference at scale. Science 380, eabn3107 (2023).

8. Rossier, V., Vesztrocy, A. W., Robinson-Rechavi, M. & Dessimoz, C. OMAmer: tree-driven and alignment-free protein assignment to subfamilies outperforms closest sequence approaches. Bioinformatics (2021) doi:10.1093/bioinformatics/btab219.

9. Steinegger, M. & Söding, J. Clustering huge protein sequence sets in linear time. Nat. Commun. 9, 2542 (2018).

10. Altenhoff, A. M. et al. Standardized benchmarking in the quest for orthologs. Nat. Methods 13, 425–430 (2016).

11. Schoch, C. L. et al. NCBI Taxonomy: a comprehensive update on curation, resources and tools. Database 2020, (2020).

12. Kumar, S. et al. TimeTree 5: An Expanded Resource for Species Divergence Times. Mol. Biol. Evol. 39, (2022).

13. Zajac, N. et al. Gene Duplication and Gain in the Trematode Atriophallophorus winterbourni Contributes to Adaptation to Parasitism. Genome Biol. Evol. 13, (2021).

## Online methods references

14. Hartuv, E. & Shamir, R. A clustering algorithm based on graph connectivity. Inf. Process. Lett. 76, 175–181 (2000).

15. Katoh, K. & Standley, D. M. MAFFT multiple sequence alignment software version 7: improvements in performance and usability. Mol. Biol. Evol. 30, 772–780 (2013).

16. Price, M. N., Dehal, P. S. & Arkin, A. P. FastTree 2--approximately maximum-likelihood trees for large alignments. PLoS One 5, e9490 (2010).

17. Yang, Z. Computational Molecular Evolution. (OUP Oxford, 2006).

18. Huerta-Cepas, J., Dopazo, H., Dopazo, J. & Gabaldón, T. The human phylome. Genome Biol. 8, R109 (2007).

19. Dylus, D., Altenhoff, A., Majidian, S., Sedlazeck, F. J. & Dessimoz, C. Inference of phylogenetic trees directly from raw sequencing reads using Read2Tree. Nat. Biotechnol. (2023) doi:10.1038/s41587-023-01753-4.

20. Dylus, D. et al. How to build phylogenetic species trees with OMA. F1000Res. 9, 511 (2020).

21. Di Tommaso, P. et al. Nextflow enables reproducible computational workflows. Nat. Biotechnol. 35, 316–319 (2017).

22. Cosentino, S. & Iwasaki, W. SonicParanoid2: fast, accurate, and comprehensive orthology inference with machine learning and language models. bioRxiv 2023.05.14.540736 (2023) doi:10.1101/2023.05.14.540736.

23. Emms, D. M. & Kelly, S. OrthoFinder: phylogenetic orthology inference for comparative genomics. Genome Biol. 20, 238 (2019).

24. Huerta-Cepas, J., Serra, F. & Bork, P. ETE 3: Reconstruction, Analysis, and Visualization of Phylogenomic Data. Mol. Biol. Evol. 33, 1635–1638 (2016).

25. Train, C.-M., Pignatelli, M., Altenhoff, A. & Dessimoz, C. iHam & pyHam: visualizing and processing hierarchical orthologous groups. Bioinformatics (2018) doi:10.1093/bioinformatics/bty994.

26. Nevers, Y. et al. Multifaceted quality assessment of gene repertoire annotation with OMArk. bioRxiv 2022.11.25.517970 (2022) doi:10.1101/2022.11.25.517970.

27. Nevers, Y. et al. The Quest for Orthologs orthology benchmark service in 2022. Nucleic Acids Res. 50, W623–W632 (2022).

28. Vilella, A. J. et al. EnsemblCompara GeneTrees: Complete, duplication-aware phylogenetic trees in vertebrates. Genome Res. 19, 327–335 (2009).

29. Persson, E., Kaduk, M., Forslund, S. K. & Sonnhammer, E. L. L. Domainoid: domain-oriented orthology inference. BMC Bioinformatics 20, 523 (2019).

30. Li, L., Stoeckert, C. J., Jr & Roos, D. S. OrthoMCL: identification of ortholog groups for eukaryotic genomes. Genome Res. 13, 2178–2189 (2003).

31. Nevers, Y. et al. OrthoInspector 3.0: open portal for comparative genomics. Nucleic Acids Res. 47, D411–D418 (2019).

32. Cosentino, S. & Iwasaki, W. SonicParanoid: fast, accurate and easy orthology inference. Bioinformatics 35, 149–151 (2019).

33. Mi, H. et al. PANTHER version 16: a revised family classification, tree-based classification tool, enhancer regions and extensive API. Nucleic Acids Res. 49, D394–D403 (2021).

34. Schreiber, F. & Sonnhammer, E. L. L. Hieranoid: hierarchical orthology inference. J. Mol. Biol. 425, 2072–2081 (2013).

35. Altenhoff, A. M. et al. OMA standalone: orthology inference among public and custom genomes and transcriptomes. Genome Res. 29, 1152–1163 (2019).

36. Altenhoff, A. M., Gil, M., Gonnet, G. H. & Dessimoz, C. Inferring hierarchical orthologous groups from orthologous gene pairs. PLoS One 8, e53786 (2013).

37. Train, C.-M., Glover, N. M., Gonnet, G. H., Altenhoff, A. M. & Dessimoz, C. Orthologous Matrix (OMA) algorithm 2.0: more robust to asymmetric evolutionary rates and more scalable hierarchical orthologous group inference. Bioinformatics 33, i75–i82 (2017).

38. Zahn-Zabal, M., Dessimoz, C. & Glover, N. M. Identifying orthologs with OMA: A primer. F1000Res. 9, 27 (2020).

39. Jones, P. et al. InterProScan 5: genome-scale protein function classification. Bioinformatics 30, 1236–1240 (2014).

40. Blum, M. et al. The InterPro protein families and domains database: 20 years on. Nucleic Acids Res. 49, D344–D354 (2021).

41. Fernández, R., Gabaldon, T. & Dessimoz, C. Orthology: Definitions, prediction, and impact on species phylogeny inference. Phylogenetics in the Genomic Era 2.4:1–2.4:14 (2020).

42. Altenhoff, A. M. & Dessimoz, C. Phylogenetic and functional assessment of orthologs inference projects and methods. PLoS Comput. Biol. 5, e1000262 (2009).

43. Boeckmann, B., Robinson-Rechavi, M., Xenarios, I. & Dessimoz, C. Conceptual framework and pilot study to benchmark phylogenomic databases based on reference gene trees. Brief. Bioinform. 12, 423–435 (2011).

44. Emms, D. M. & Kelly, S. Benchmarking Orthogroup Inference Accuracy: Revisiting Orthobench. Genome Biol. Evol. 12, 2258–2266 (2020).

